# Balanced Input from the tRNA Prenyltransferase MiaA Controls the Stress Resistance and Virulence Potential of Extraintestinal Pathogenic *Escherichia coli*

**DOI:** 10.1101/2021.02.02.429414

**Authors:** Matthew G. Blango, Brittany A. Fleming, William M. Kincannon, Alex Tran, Adam J. Lewis, Colin W. Russell, Qin Zhou, Lisa M. Baird, John R. Brannon, Connor J. Beebout, Vahe Bandarian, Maria Hadjifraniskou, Michael T. Howard, Matthew A. Mulvey

## Abstract

An ability to adapt to rapidly changing and often hostile environments is key to the success of many bacterial pathogens. In *Escherichia coli*, the highly conserved enzymes MiaA and MiaB mediate the sequential prenylation and methylthiolation of adenosine-37 within tRNAs that decode UNN codons. Here, we show that MiaA, but not MiaB, is critical to the fitness and virulence of extraintestinal pathogenic *E. coli* (ExPEC), a major cause of urinary tract and bloodstream infections. Deletion of *miaA* has pleiotropic effects, rendering ExPEC especially sensitive to stressors like nitrogen and oxygen radicals and osmotic shock. We find that stress can stimulate striking changes in *miaA* expression, which in turn can increase translational frameshifting and markedly alter the bacterial proteome. Cumulatively, these data indicate that ExPEC, and likely other organisms, can vary MiaA levels as a means to fine-tune translation and the spectrum of expressed proteins in response to changing environmental challenges.

## INTRODUCTION

The translation of mRNA into protein by ribosomes and aminoacyl-transfer RNA (tRNA) complexes is an energy-intensive process that is subject to multiple levels of complicated regulation. For example, tRNAs can be covalently modified by more than 100 different moieties that can influence the charging of tRNAs with amino acids, tRNA stability, codon usage, and reading frame maintenance [1–4]. In *Escherichia coli* and other bacteria, the hypomodification of tRNAs can result in decreased growth rates, altered metabolic requirements, and reduced stress resistance [5–8]. Loss of tRNA modifications can also impact the fitness and virulence potential of many important bacterial pathogens, including *Streptomyces pyogenes*, *Pseudomonas spp.*, *Shigella flexneri*, *Agrobacterium tumefaciens, Mycobacterium tuberculosis, Aeromonas hydrophila*, *Streptococcus spp.*, and *Salmonella enterica* serotype Typhimurium [6, 9–21]. Together, these findings suggest that tRNA modification serves as a regulatory nexus that can control a wide array of bacterial activities.

One of the most commonly modified tRNA residues in bacteria is adenosine-37 (A-37), which lies adjacent to the anticodon loop [8, 22]. In its final form in *E. coli*, A-37 of UNN-recognizing tRNA molecules is oftentimes prenylated and methylthiolated [23]. The *miaA* gene of *E. coli* encodes a tRNA prenyltransferase that catalyzes the addition of a prenyl group onto the *N^6^*-nitrogen of A-37 to create i^6^A-37 tRNA [24, 25] (Fig. 1A). The modified i^6^A-37 residue is subsequently methylthiolated by the radical-S-adenosylmethionine enzyme MiaB to create ms^2^i^6^A-37 [26]. The bulky and hydrophobic ms^2^i^6^A-37 modification enhances tRNA interactions with UNN target codons, promoting reading frame maintenance and translational fidelity [5, 8, 27]. Mutations in the *miaA* locus result in an unmodified A-37 residue, as prenylation is required for methylthiolation by MiaB. In K-12 laboratory-adpated *E. coli* strains, mutations in *miaA* impair attenuation of the tryptophan and phenylalanine operons [28, 29] and diminish translation of the stationary phase sigma factor RpoS and the small RNA chaperone Hfq [7, 30, 31]. Additionally, mutants lacking *miaA* are unable to effectively resolve aberrant DNA-protein crosslinks [32] and have somewhat elevated spontaneous mutation frequencies [33–35]. The ms^2^i^6^A-37 modification is highly conserved in both prokaryotes and eukaryotes, though the specific enzymes that mediate this modification have diverged within evolutionarily distant organisms [8]. However, in prokaryotes, MiaA and MiaB homologues are relatively well conserved, and the enzymes appear to function similarly in all tested bacterial species [36, 37].

**Figure 1.**
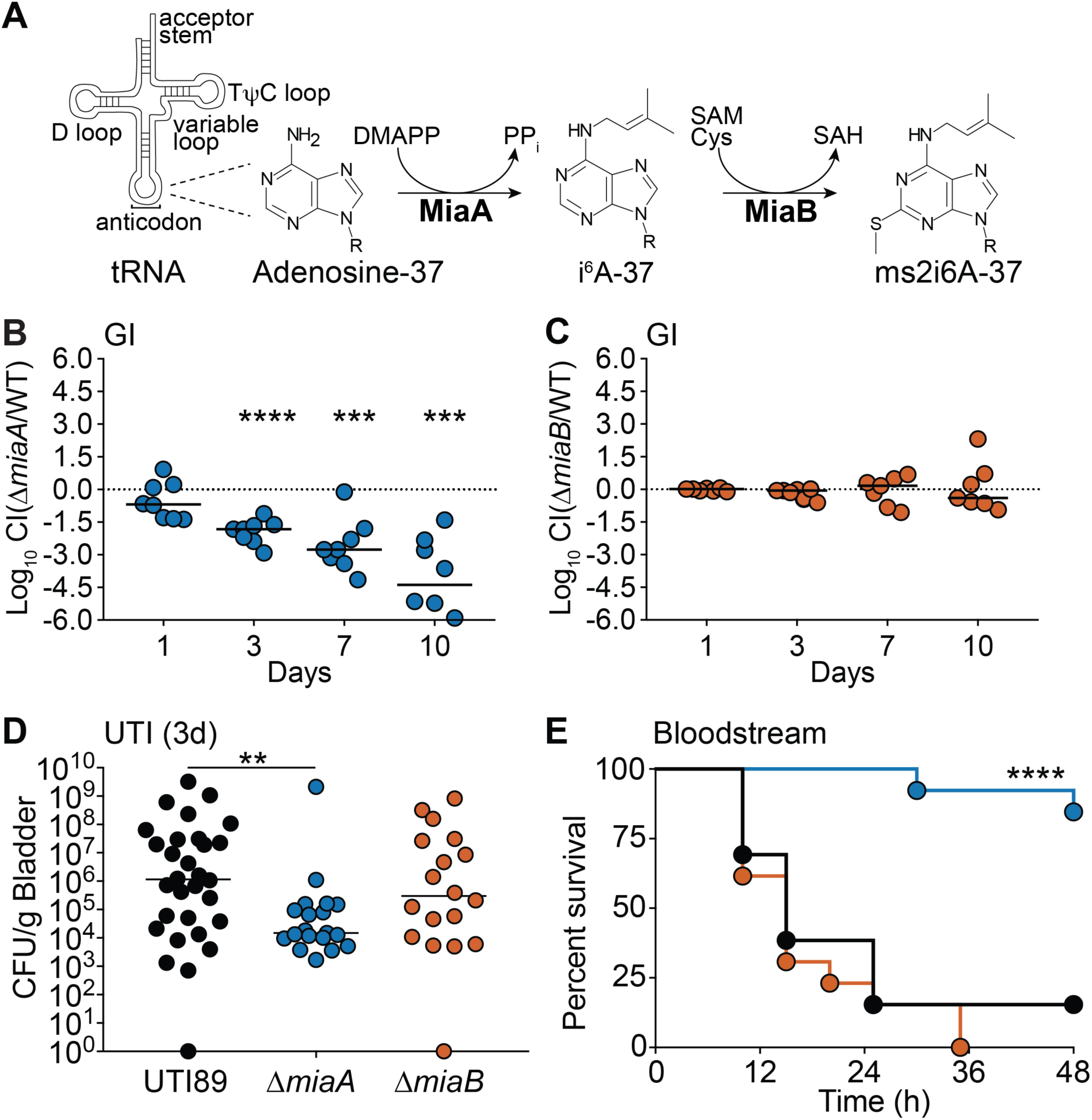
MiaA promotes ExPEC fitness and virulence within diverse host niches. (**A**) MiaA and MiaB act sequentially to modify tRNA molecules that recognize UNN codons; modified from [37]. DMAPP, dimethylallyl diphosphate; SAM, S-adenosylmethionine; SAH, S-adenosylhomocysteine; Cys, cysteine. (**B** and **C**) To assess gut colonization, adult BALB/c mice were inoculated via oral gavage with ∼10^9^ CFU of a 1:1 mixture of (**B**) UTI89 and UTI89Δ*miaA* or (**C**) UTI89 and UTI89Δ*miaB.* Fecal titers were determined at the indicated time points and used to calculate competitive indices (CI). ***, *P* < 0.001; ****, *P* < 0.0001 by one sample *t*-tests. *n* = 7-8 mice from two independent experiments. (**D**) The bladders of adult female CBA/J mice were inoculated via transurethral catheterization with ∼10^7^ CFU of UTI89, UTI89Δ*miaA*, or UTI89Δ*miaB*. Mice were sacrificed 3 days later and bacterial titers within the bladders were determined by plating tissue homogenates. **, *P* < 0.01 by Mann Whitney U tests; *n* ≥ 19 mice per group from at least three independent experiments. In B, C, and D, bars indicate median values; dots represent individual mice. (**E**) Kaplan Meier survival curves of C57Bl/6 mice inoculated via i.p. injections with ∼10^7^ CFU of UTI89 (black line), UTI89Δ*miaA* (blue), or UTI89Δ*miaB* (orange). ****, *P* < 0.0001 by Log-rank Mantel Cox test for UTI89 versus UTI89Δ*miaA*; *n* = 13 mice per group from two independent experiments.

Given that the ms^2^i^6^A-37 modification is a well-defined regulator of many tRNA functions in lab-adapted K-12 *E. coli* strains, we sought to understand how this modification is co-opted in a pathogenic *E. coli* background. *E. coli* pathotypes display extensive genetic diversity and are usually more resilient under stress than their lab-adapted counterparts [38]. Extraintestinal Pathogenic *E. coli* (ExPEC) typically reside in the lower intestinal tract of mammals, where they are rarely associated with pathology [39]. However, when they spread outside the gut to other host sites ExPEC can cause a number of serious diseases, including urinary tract and bloodstream infections [38, 40]. Bacterial pathogens like ExPEC must be able to rapidly respond to a diverse array of stressors encountered within changing host environments. These include nutrient deprivation, redox stress in the form of oxygen and nitrogen radicals, extremes in pH, envelope damage, changing osmotic pressures, and a wide assortment of host immune effector cells and antimicrobial compounds [41–45].

Shifts in the prevalence of specific tRNA modifications, such as A-37 prenylation mediated by MiaA, are proposed to help optimize bacterial responses to stress by affecting translational fidelity and selective protein expression [14, 33, 46]. In other words, changing levels of tRNA modifications may control the codon-biased translation of select transcripts, providing a post-transcriptional programmable mechanism that distressed cells can use to facilitate beneficial changes in their proteomes. Here we provide evidence in support of this hypothesis, showing that ExPEC can modulate MiaA levels in response to stress, and that varying levels of this enzyme can increase translational frameshifting and markedly alter the spectrum of expressed proteins. Furthermore, our data reveal that MiaA, but not MiaB, is critical to the fitness and virulence of ExPEC in both *in vitro* assays and in mouse models of infection and intestinal colonization.

## RESULTS

### MiaA promotes ExPEC fitness and virulence in vivo

To assess the importance of MiaA and MiaB for ExPEC within varied host environments, we employed well-established mouse models of gut colonization, urinary tract infection (UTI), and bloodstream infection [47]. For these and subsequent experiments, *miaA* and *miaB* were independently deleted from the ExPEC reference strain UTI89 to generate the isogenic knockout mutants UTI89Δ*miaA* and UTI89Δ*miaB* [48, 49].

#### Gut colonization

The mammalian gastrointestinal (GI) tract serves as a major reservoir for ExPEC that can seed extraintestinal infections [50–54]. Roles for MiaA and MiaB in ExPEC colonization of the GI tract were probed using competitive assays in which ∼10^9^ colony forming units (CFU) of a 1:1 mixture of UTI89 and either UTI89Δ*miaA* or UTI89Δ*miaB* were introduced into adult specific-pathogen-free (SPF) BALB/c mice via intragastric gavage [55–57]. In this model system, the levels of ExPEC recovered from the feces reflect ExPEC titers within the large intestines [55]. For these assays, UTI89 and the *miaA* and *miaB* knockout mutants were engineered to express either kanamycin (Kan^R^) or chloramphenicol (Cam^R^) resistance cassettes so that the strains could be readily identified by plating fecal homogenates on selective media. Feces were collected at the indicated time points and the numbers of viable bacteria were enumerated to determine competitive indices (CI). UTI89Δ*miaA* was significantly outcompeted by wild-type UTI89 as early as day 3 post-inoculation (Fig. 1B). By day 10, there was about a 25,000-fold reduction in the relative levels of UTI89Δ*miaA* recovered from the feces, correlating with a median CI of −4.39. At this time point, UTI89Δ*miaA* titers in the majority of mice were below the limit of detection. In contrast, there were no notable differences in titers between UTI89Δ*miaB* and UTI89 in the feces at any time point (Fig. 1C). These results indicate that the loss of MiaA, but not MiaB, greatly impairs the fitness of UTI89 within the gut

#### UTI

During the course of a UTI, ExPEC is able to bind and invade the host epithelial cells that comprise the bladder mucosa [58]. Once internalized into bladder cells, ExPEC can traffic into late endosome-like compartments where it may form quiescent reservoir populations that promote long-term bacterial persistence. Alternatively, ExPEC can enter the host cytosol and rapidly multiply, forming large intracellular bacterial communities that eventually rupture the epithelial cell. In cell culture-based assays using a bladder epithelial cell line, we found that UTI89Δ*miaA* and UTI89Δ*miaB* are able to bind, invade, and survive intracellularly in overnight assays much like wild-type UTI89 (Supplemental Fig. S1).

To investigate MiaA and MiaB requirements during UTI, 10^7^ CFU of wild-type UTI89, UTI89Δ*miaA*, and UTI89Δ*miaB* were independently inoculated via transurethral catheterization into adult female CBA/J mice and bacterial titers in the bladders were determined after 3 days. In this analysis, UTI89Δ*miaB* showed no statistically significant defect relative to the parent strain; whereas the Δ*miaA* strain was clearly attenuated (Fig. 1D). Deficiencies in bladder colonization by UTI89Δ*miaA* were apparent by 6 h post-inoculation (Supplemental Fig. S2A), and were still significant after 9 days (Supplemental Fig. S2B). The differences observed between wild-type UTI89 and UTI89Δ*miaA* at 3 days post-inoculation of CBA/J mice were also manifest in C3H/HeJ mice (Supplemental Fig. S2C). Due to defects in Toll-like receptor 4 (TLR4) signaling and other innate defenses, C3H/HeJ mice have attenuated inflammatory responses and increased susceptibility to UTI [59–62]. Our results indicate that the decreased capacity of UTI89Δ*miaA* to colonize the bladder is not attributable to an inability of the *miaA* knockout to handle TLR4-dependent innate host defenses. Collectively, these results indicate that MiaA is required for maximal fitness in mouse UTI models, while MiaB is less critical.

#### Bloodstream infection

ExPEC is a leading cause of bloodstream infections, which too often trigger discordant systemic inflammatory responses that can result in a life-threatening condition known as sepsis [63]. To examine the contributions of MiaA and MiaB to ExPEC virulence and fitness in a model of sepsis, adult C57Bl/6 mice were inoculated via intraperitoneal (i.p.) injections with ∼10^7^ CFU of wild-type UTI89, UTI89Δ*miaA*, or UTI89Δ*miaB.* Following i.p. injection, the bacteria enter the bloodstream and disseminate [56, 64, 65]. In our experiments, only 15% (2/13) of the mice infected with wild-type UTI89, and 0% (0/13) of the mice injected with UTI89Δ*miaB,* were viable after 48 hours (Fig. 1E). In sharp contrast, 84% (11/13) of the mice infected with UTI89Δ*miaA* survived. At six hours post-injection, significantly lower numbers of bacteria were recovered from the spleens and kidneys of UTI89Δ*miaA*-infected mice, relative to mice infected with wild-type UTI89 or UTI89Δ*miaB* (Supplemental Fig. S3A and B). While not significant, titers in the liver also trended lower in UTI89Δ*miaA*-infected mice (Supplemental Fig. S3C). Combined, these data demonstrate that MiaA is important for the virulence of ExPEC and its survival during systemic infections, while MiaB appears dispensable.

### MiaA enhances ExPEC growth and stress resistance

Earlier studies showed that K-12 *E. coli* and *Salmonella* mutants lacking *miaA* are moderately impaired in nutrient-rich broth, but less so in nutrient-limited media [27, 35, 66]. Using *in vitro* growth assays, we found that UTI89Δ*miaA* grew normally in modified M9 minimal media, but failed to reach densities as high as the wild-type strain in more complex, nutrient-rich lysogeny broth (LB) (Fig. 2A **and** B). In contrast, the *miaB* knockout exhibited no overt growth defects in either type of media. These data suggest that UTI89Δ*miaA* has reduced metabolic flexibility relative to wild-type UTI89 and the *miaB* mutant. This may contribute to the decreased fitness of UTI89Δ*miaA* in our mouse models, where the bacteria likely encounter marked shifts in nutrient availability. However, within different host environments ExPEC will face a wide variety of additional challenges that might be countered by MiaA-dependent processes. We investigated this possibility by examining the effects of MiaA and MiaB on ExPEC resistance to nitrosative, oxidative, and osmotic stress.

**Figure 2.**
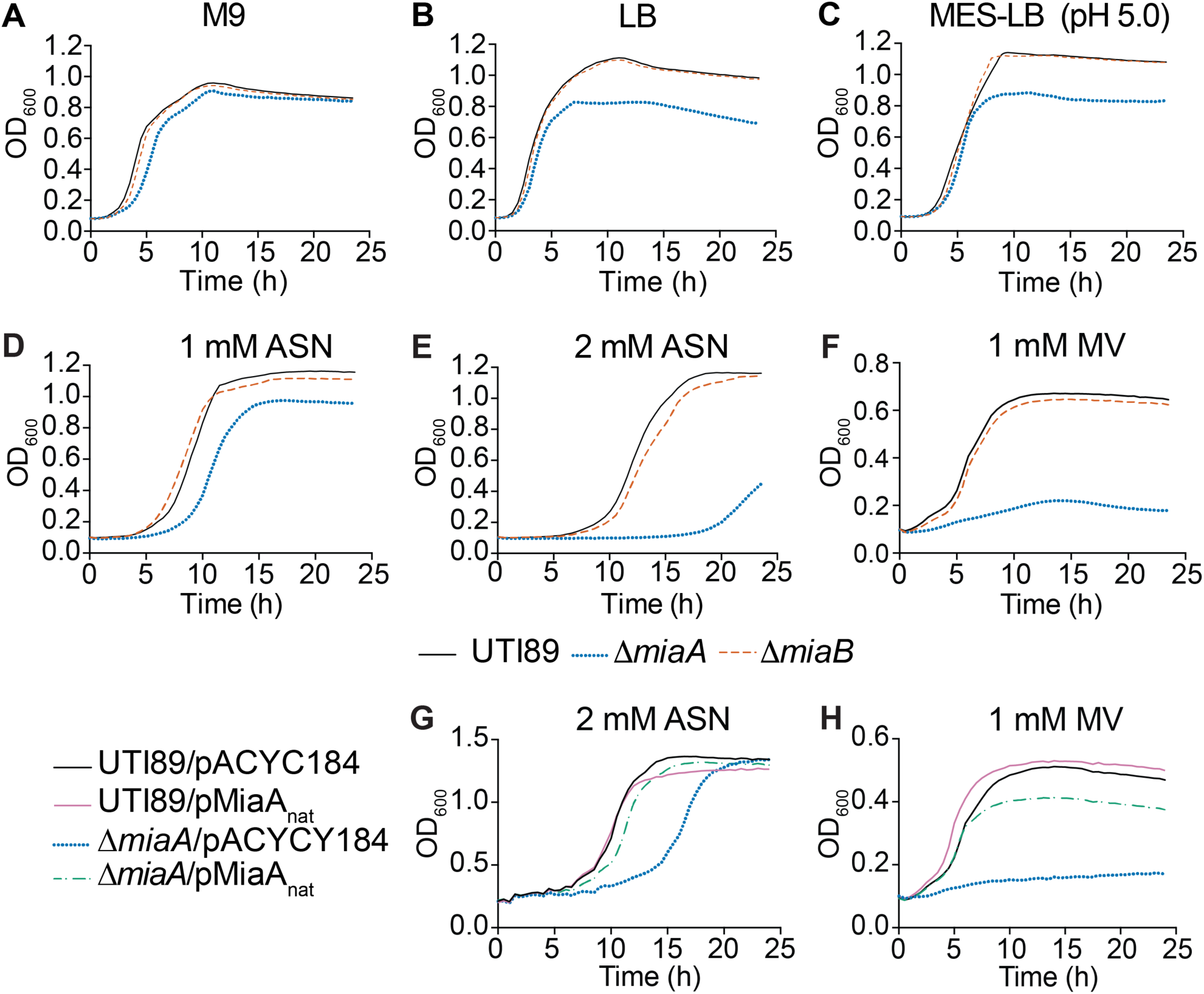
Deletion of *miaA* limits growth of UTI89 in rich medium and lowers resistance to oxidative and nitrosative stress. (**A-F**) Graphs indicate mean growth of UTI89, UTI89Δ*miaA*, and UTI89Δ*miaB* in shaking cultures with modified M9 media, LB, MES-LB, MES-LB with 1 or 2 mM ASN, or LB with 1 mM MV. (**G** and **H**) Curves show mean growth of UTI89 and UTI89Δ*miaA* carrying pMiaA_nat_ (with *miaA* expressed from its native promoter) or the empty vector control pACYC184 in LB containing (G) 2 mM ASN or (H) 1 mM MV. Each curve shows the means of results from four replicates, and are representative of three independent experiments.

#### Oxidative and nitrosative stress

During the course of an infection, both host and bacterial cells can produce reactive oxygen and nitrogen radicals that can damage lipids, proteins, and nucleic acids [67, 68]. The contributions of MiaA and MiaB to nitrosative and oxidative stress resistance were assessed using acidified sodium nitrite (ASN) and methyl viologen (MV), respectively. When added to low pH morpholineethanesulfonic acid (MES)-buffered LB (MES-LB; pH 5.0), sodium nitrite dismutates to form nitrous acid which in turn generates NO and other harmful reactive nitrogen species [69]. In un-supplemented MES-LB, UTI89Δ*miaA* reached a lower maximal density than wild-type UTI89 (Fig. 2C), similar to results obtained using standard LB (Fig. 2B). The addition 1 mM ASN delayed entry of UTI89Δ*miaA* into exponential growth phase by close to 4 hours (Fig. 2D), while 2 mM ASN delayed growth by more than 15 hours relative to wild-type UTI89 (Fig. 2E). The addition of 1 mM MV, which produces superoxide radicals [70], had even stronger inhibitory effects on growth of UTI89Δ*miaA* (Fig. 2F). In contrast, UTI89Δ*miaB* grew much like the wild-type strain in the presence of ASN or MV (Fig. 2D-F). Complementation with pMiaA_nat_, a low copy plasmid that encodes MiaA under control of its native promoter, restored growth of UTI89Δ*miaA* to near wild-type levels in both 2 mM ASN (Fig. 2G) and in 1 mM MV (Fig. 2H).

#### Osmotic stress

During a UTI, osmotic pressure within the bladder can shift from 50 to >1,400 mOsm/kg due to varying concentrations of solutes like sodium and urea [71, 72]. By comparison, the normal osmolarity of blood ranges from about 275 to 295 mOsm/kg. To test the sensitivities of wild-type UTI89 and the knockout strains to hypoosmotic stress, we diluted the bacteria from early stationary phase cultures into ddH_2_O, and then quantified the number of viable bacteria every 30 minutes over the course of 2 hours. Titers of UTI89*ΔmiaA* carrying the empty vector pACYC184 were greatly reduced following exposure to hypoosmotic stress, whereas the levels of UTI89/pACYC184 and UTI89Δ*miaB*/pACYC184 remained mostly unchanged (Fig. 3A). Survival of UTI89Δ*miaA* was restored by complementation with pMiaA_nat_. To ensure that reduced survival of UTI89Δ*miaA* was attributable to hypoosmotic stress and not starvation, cells were resuspended in ddH_2_O containing 0.1% glucose, which is comparable to the glucose levels within our M9 medium. Viable bacteria measured after 120 minutes indicated that the death of UTI89Δ*miaA* was not due to nutrient deprivation (Fig. 3B). We also observed that UTI89Δ*miaA* grew poorly in hyperosmotic conditions, created by addition of 5% NaCl to standard LB (Fig. 3C). As in other assays, UTI89Δ*miaB* behaved more like the wild-type strain. Growth of UTI89Δ*miaA* was restored to wild-type levels by complementation with pMiaA_nat_ (Fig. 3D). These results indicate that the *miaA* knockout has decreased resistance to both hypo- and hyperosmotic stresses.

**Figure 3.**
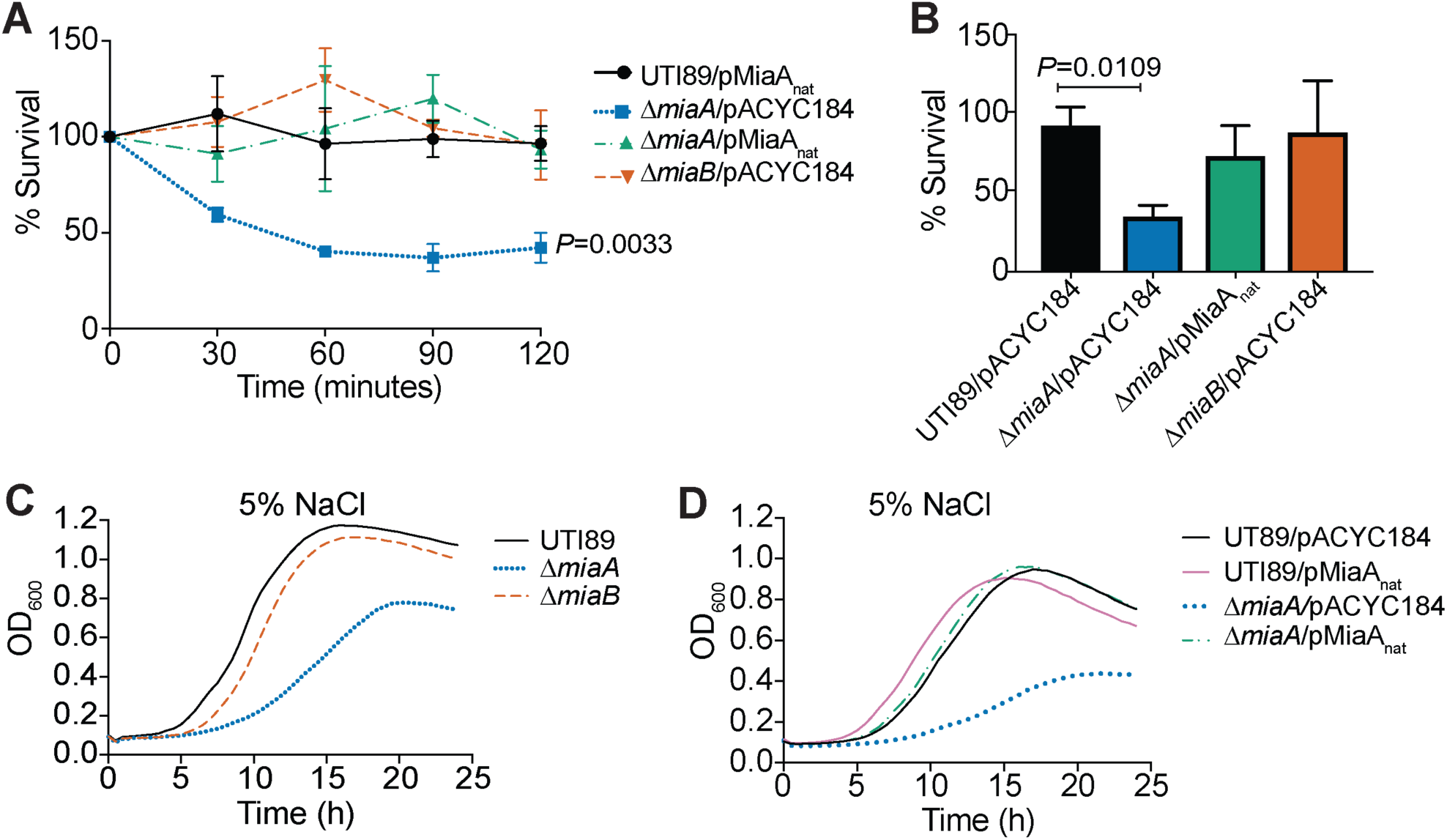
MiaA enhances the resistance of UTI89 to osmotic stress. (**A**) Bacteria were grown to stationary phase in LB, pelleted, and resuspended in ddH_2_O. The mean numbers (±SEM) of surviving bacteria recovered at the indicated time points are graphed as the percentage of the bacteria present immediately after resuspension in ddH_2_O (time 0). *P*-values were determined by two-way ANOVA with the Geisser-Greenhouse correction; n = 3 independent assays each done in triplicate. (**B**) Bars indicate mean numbers of bacteria (±SD) that survived 2 hours in ddH_2_O with 0.1% glucose, calculated as a percent of the inoculum. *P* values were determined, relative to the control strain UTI89/pMiaA_nat_, by unpaired *t*-tests with Welch’s correction; n = 3 independent assays. (**C** and **D**) Curves show growth of the indicated strains in LB plus 5% NaCl, as measured by OD_600_. Data are representative of at least three independent experiments performed in quadruplicate.

### Hyperosmotic stress attenuates MiaA translation

We next examined how MiaA levels in UTI89 change in response to environmental cues, focusing on hyperosmotic stress. For these assays, we employed a low-copy number plasmid (pMiaA-Flag_nat_) that encodes C-terminal FLAG-tagged MiaA under control of the native *miaA* promoter. Mid-logarithmic phase cultures of UTI89/pMiaA-Flag_nat_ were resuspended in LB ± 5% NaCl and levels of MiaA-Flag were then assessed by western blot at 30-minute intervals over the course of 1.5 hours (Fig. 4A). Interestingly, MiaA levels in UTI89 exposed to high salt broth were decreased at all time points in comparison with bacteria grown in standard LB. We observed a similar phenomenon if overnight cultures of UTI89/pMiaA-Flag_nat_ in standard LB were back-diluted into high salt broth and then grown to mid-logarithmic phase (OD_600_≍0.5, Fig. 4B). Of note, in these assays we observed no loss of the pMiaA-Flag_nat_ construct. In addition, *miaA* transcripts were often elevated following exposure of UTI89 to high salt (Fig. 4C), suggesting that the downregulation of MiaA protein levels in response to this osmotic stress occurs via a post-transcriptional mechanism. The transcription of *miaB* was notably reduced under the same conditions (Fig. 4D).

**Figure 4.**
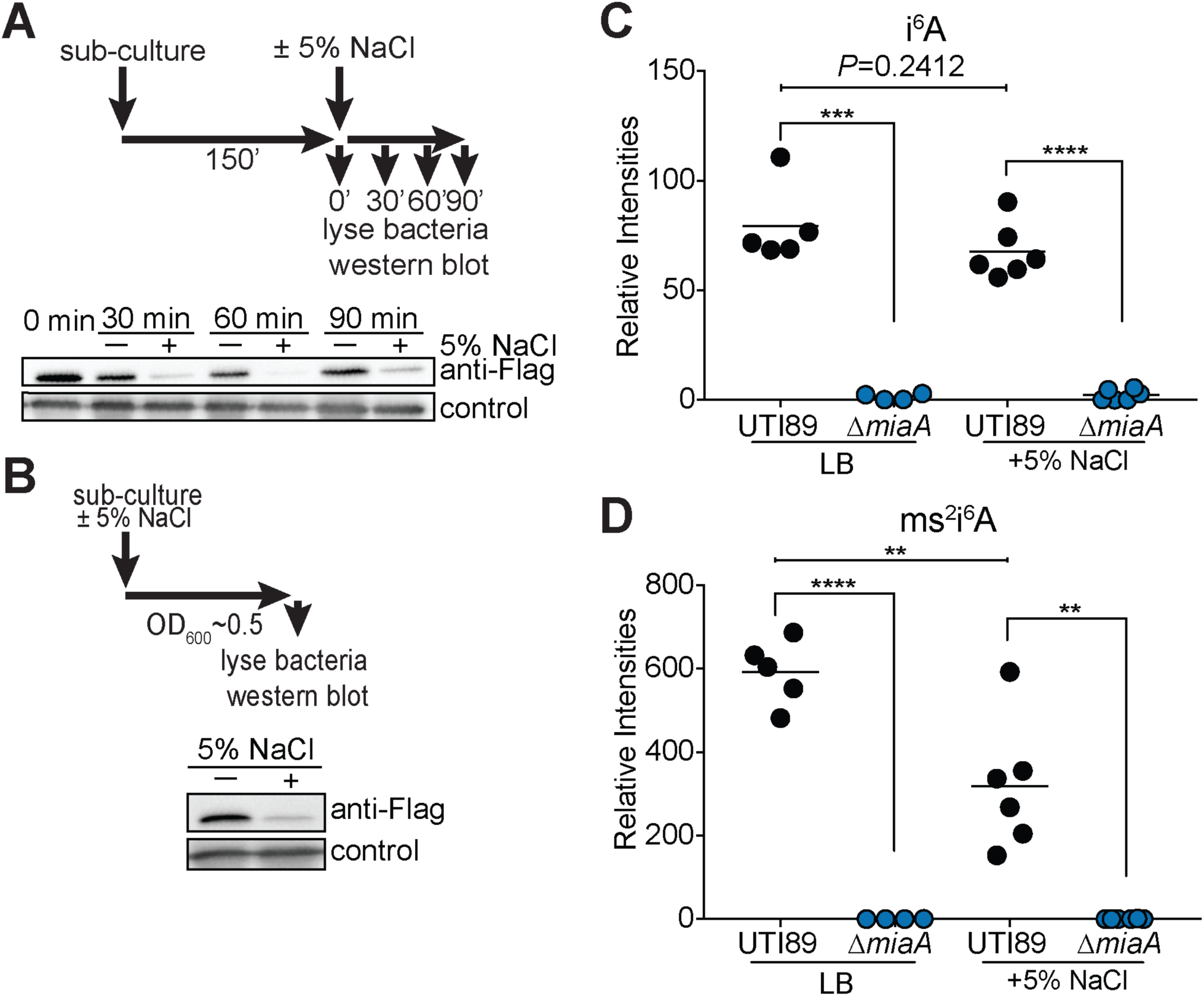
High salt stress downregulates MiaA translation and reduces ms^2^i^6^A levels. (**A**) Top panel shows schematic of the experimental setup. UTI89/pMiaA-Flag_nat_ was diluted from overnight cultures into fresh LB and grown shaking for 2.5 hours at 37°C prior to resuspension in LB or LB + 5% NaCl. Incubations were continued for the indicated times before samples were collected and analyzed by western blots using anti-Flag and anti-*E. coli* (loading control) antibodies (bottom panel). (**B**) Top panel summarizes the experimental setup. Overnight cultures of UTI89/pMiaA-Flag_nat_ were diluted directly into fresh LB or LB + 5% NaCl and grown shaking to OD_600_ of 0.5 prior to processing for western blot analysis (bottom panel). (**C and D**) UTI89 from mid-logarithmic cultures in LB was resuspended in LB or LB + 5% NaCl and one hour later the evels of *miaA* and *miaB* transcripts were determined by RT-qPCR. Bars indicate mean values from 9 independent replicates, each with two technical replicates. *, *P* < 0.05; ***, *P* < 0.001 by Mann-Whitney U tests. (**E and F**) Graphs show relative levels of i^6^A and ms^2^i^6^A recovered from UTI89 and UTI89Δ*miaA* following growth to OD_600_ of 0.5 in LB or LB + 5% NaCl, as determined by LC-MS. **, *P* < 0.01; ***, *P* < 0.001; ****, *P* < 0.0001 by unpaired *t* tests. Bars indicate median values from 4 to 6 independent replicates.

To determine if lower amounts of the MiaA protein detected in high salt broth culture affected i^6^A or ms^2^i^6^A levels, we employed liquid chromatography-coupled mass spectrometry (LC-MS). Normalized amounts of the i^6^A modification in wild-type UT89 grown to mid-logarithmic phase in LB were similar to those measured in UTI89 grown in high salt broth (Fig. 4E). However, hyperosmotic stress caused a marked reduction in relative ms^2^i^6^A levels (Fig. 4F), possibly due to reduced transcription of *miaB* (Fig. 4D). i^6^A and ms^2^i^6^A were undetectable in UTI89Δ*miaA*, regardless of high salt exposure, confirming that MiaA is required for both modifications (Fig. 4E-F). In contrast, deletion of *miaB* prevented formation of ms^2^i^6^A, but led to greatly elevated levels of i^6^A (Supplemental Fig. S4). Cumulatively, these data indicate that in response to hyperosmotic stress UTI89 can post-transcriptionally downregulate MiaA, coordinate with reduction of both *miaB* messages and ms^2^i^6^A levels.

### Overexpression of MiaA is detrimental under stressful conditions

Since it was unexpected that high salt stress would lead to a decrease in the levels of MiaA and the ms^2^i^6^A modification, we set out to determine if overexpression of MiaA would affect bacterial growth during environmental stress. To overexpress MiaA, we utilized a plasmid (pRR48) with *miaA* under control of an IPTG-inducible P*tac* promoter in wild-type UTI89. By LC-MS, relative intensities of i^6^A were significantly higher in UTI89/pMiaA_P*tac*_ induced with 1 mM IPTG and grown to mid-logarithmic phase in LB compared to UTI89 carrying the empty vector pRR48, whereas the relative intensities of ms^2^i^6^A were only modestly elevated (Fig. 5A).

**Figure 5.**
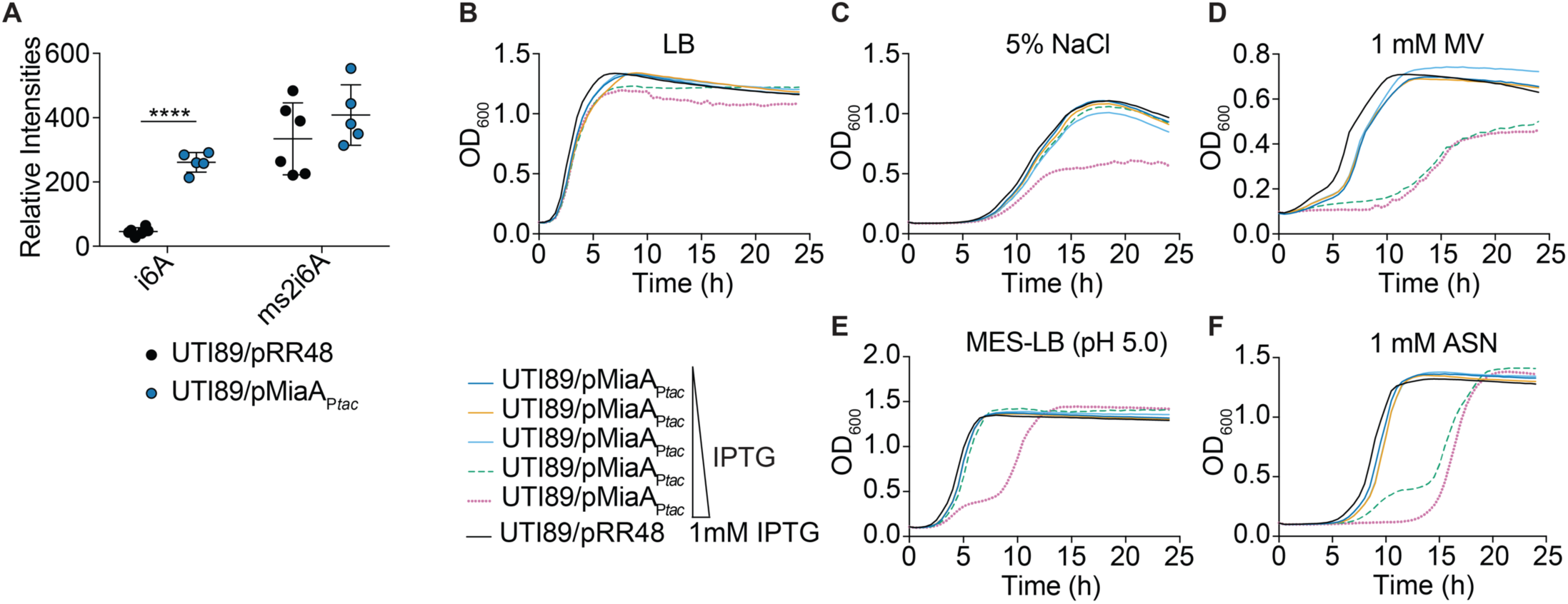
Overexpression of MiaA reduces ExPEC stress resistance. (**A**) Relative levels of i^6^A and ms^2^i^6^A in UTI89 carrying either pMiaA_P*tac*_ or the empty vector control pRR48 following growth to OD_600_ of 0.5 in LB with 1 mM IPTG, as quantified by LC-MS. ****, *P* < 0.0001 by unpaired *t* test; n = 5 to 6 independent replicates per group. (**B-F**) Curves depict growth of UTI89 carrying pMiaA_P*tac*_ or the empty vector control pRR48 in LB, LB + 5% NaCl, LB + 1mM MV, MES-LB, or MES-LB + 1 mM ASN. Cultures were grown shaking at 37°C with IPTG added in ten-fold increments from 0 to 1000 μM, as indicated. Each curve depicts the means of results from a single experiment and is representative of at least three independent experiments carried out in quadruplicate.

Next, overnight cultures of UTI89/pMiaA_P*tac*_ were back-diluted into LB, LB + 1 mM MV, LB + 5% NaCl, MES-LB, or MES-LB + 1 mM ASN, and grown in the presence of increasing IPTG concentrations (Fig. 5B-F). Lower levels of MiaA protein induction caused no overt defects and the bacteria grew much like UTI89/pRR48. However, higher levels of IPTG-induced MiaA expression hindered growth of UTI89/pMiaA_P*tac*_ in the presence of 1 mM MV, 5% NaCl, MES-LB, and 1 mM ASN. In contrast, over expression of MiaB did not affect bacteria growth in these *in vitro* assays (Supplemental Fig. S5). These findings indicate that too much MiaA can be detrimental to bacterial fitness, similar to the complete absence of the enzyme.

### Both Deletion and Overexpression of MiaA Increase Frameshifting

Previous research in K-12 *E. coli* and *Salmonella* showed that deletion of *miaA* can compromise translational fidelity, resulting in increased ribosomal frameshifting [73, 74]. To determine the effects of MiaA on frameshifting in UTI89, we utilized dual-luciferase reporter plasmids that consist of a translational fusion of firefly luciferase downstream of renilla luciferase. Linker sequences, derived from either Antizyme 1 (Az1) or HIV *gag-pol*, were placed between the two luciferase genes (Fig. 6A). The Az1-derived linker sequence contains a stop codon positioned in-frame so that a +1 frameshift must occur for read-through expression of firefly luciferase [75]. In contrast, a −1 frameshift is required for expression of firefly luciferase downstream of the HIV-derived linker [76]. Importantly, upstream of the in-frame stop codons in both linkers are UNN codons that can be recognized by MiaA-modified tRNAs. The firefly and renilla luciferases act on distinct substrates, which are used to sequentially assess levels of expression of each enzyme [75, 76]. Control plasmids in which the two luciferases are in-frame were used to normalize the data by accounting for ribosome drop-off.

**Figure 6.**
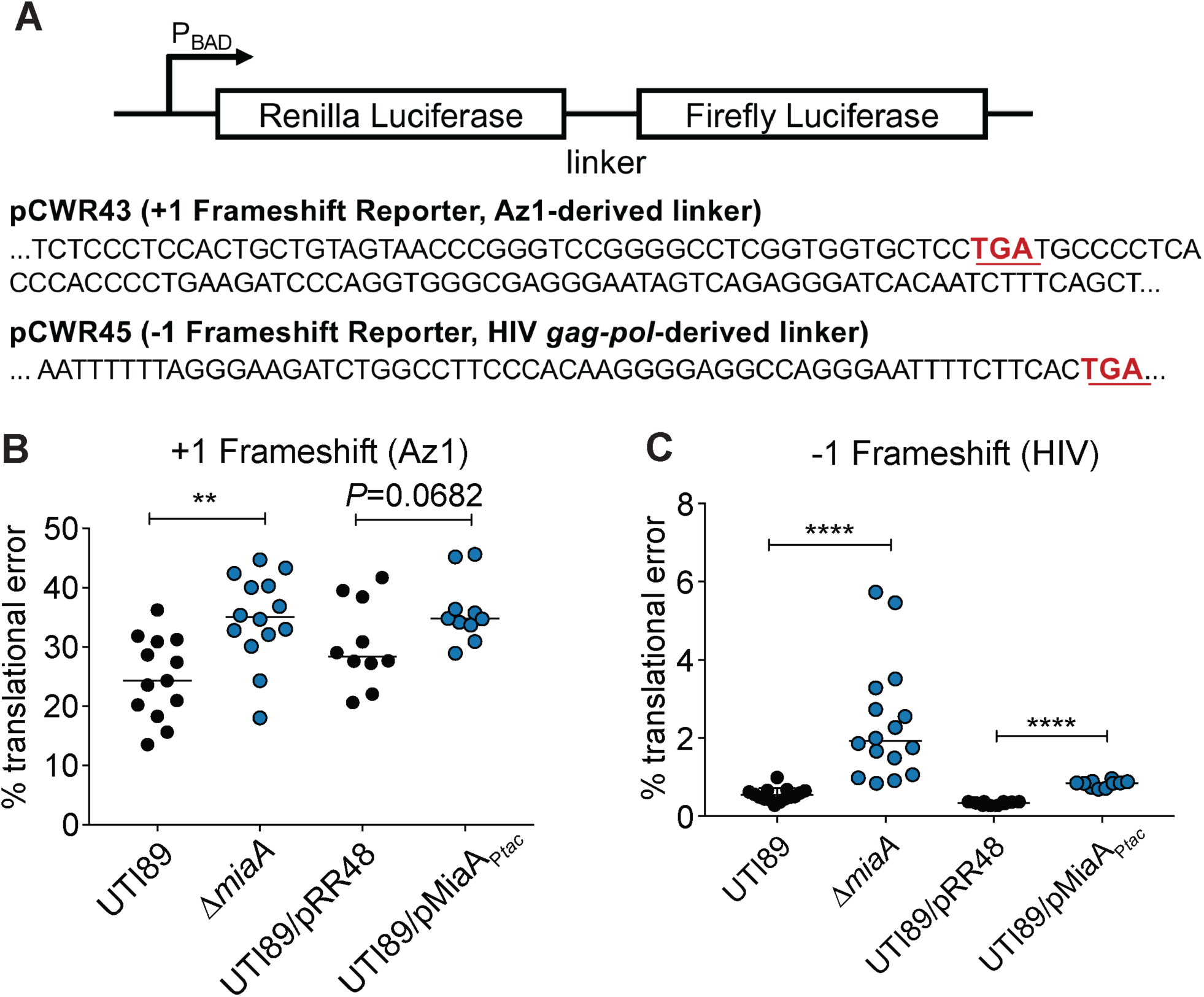
Changing levels of MiaA increase frameshifting. (**A**) Diagram depicts the structures of the dual luciferase reporters, with specific intergenic linker sequences and premature stop codons (underlined red) indicated below. Each linker contains MiaA-sensitive UNN codons. (**B and C**) Graphs show results from +1 and −1 frameshifting assays with UTI89, UTI89Δ*miaA*, UTI89/pRR48, or UTI89/pMiaA_P*tac*_ carrying one of the dual luciferase reporter constructs. Bacteria were grown shaking at 37°C in LB, with 1 mM IPTG included for UTI89/pRR48 and UTI89/pMiaA_P*tac*_. After reaching an OD_600_ of ∼0.2, 0.2% arabinose was added to induce expression of the luciferases. At an OD_600_ of 0.5, translational error rates were quantified by determining the ratio of firefly to renilla luciferase activities in bacteria carrying the +1 (Az1) and −1 (HIV) reporter constructs. Results were normalized using the ratio of firefly to renilla luciferase activity in bacteria carrying control plasmids in which the luciferases are in-frame. **, *P* < 0.01; ****, *P* < 0.0001 by two-tailed unpaired *t* tests; *n* = 10-14 independent replicates.

To examine the consequences of MiaA expression on frameshifting, the dual-luciferase reporter constructs were used in combination with wild-type UTI89, UTI89Δ*miaA*, UTI89/pMiaA_P*tac*_, and UTI89 carrying the empty control vector pRR48. After overnight growth in LB, UTI89 and UTI89Δ*miaA* were back-diluted into LB while UTI89/pMiaA_P*tac*_ and UTI89/pRR48 were back-diluted into LB + 1 mM IPTG to induce MiaA expession. After reaching mid-log growth, the enzymatic activities of the two luciferases were quantified. Both the lack of MiaA and MiaA overexpression caused notable increases in frameshifting in both the +1 and −1 directions (Fig. 6B-C). These results confirm that loss of MiaA can increase frameshifting and show that elevated MiaA levels can likewise impact the fidelity of translation.

### Changing levels of MiaA alters the spectrum of expressed proteins

To determine how deletion and overexpression of MiaA affect translation we used multidimensional protein identification technology (MudPIT; LC-MS/MS) with wild-type UTI89 and UTI89Δ*miaA* cultures grown to mid-log phase in LB, and UTI89/pMiaA_P*tac*_ and UTI89/pRR48 similarly grown in LB + 1mM IPTG. Of 1,524 proteins detected in UTI89 and UTI89Δ*miaA*, 105 were picked up only in the wild-type strain and 23 were unique to the *miaA* knockout mutant (Fig. 7A). 1,471 proteins were identified in UTI89/pRR48 and UTI89/pMiaA_P*tac*_, with 42 being exclusive to UTI89/pRR48 and 20 seen only in the MiaA overexpression strain (Fig. 7B). 115 proteins were significantly downregulated in UTI89Δ*miaA* relative to wild-type UTI89, while 34 proteins were upregulated in the knockout mutant (Fig. 7C). Notably fewer proteins were significantly altered when MiaA was overexpressed. Relative to the control strain UTI89/pRR48, 20 proteins were downregulated in UTI89/pMiaA_P*tac*_, whereas nine (including MiaA) were upregulated (Fig. 7D).

**Figure 7.**
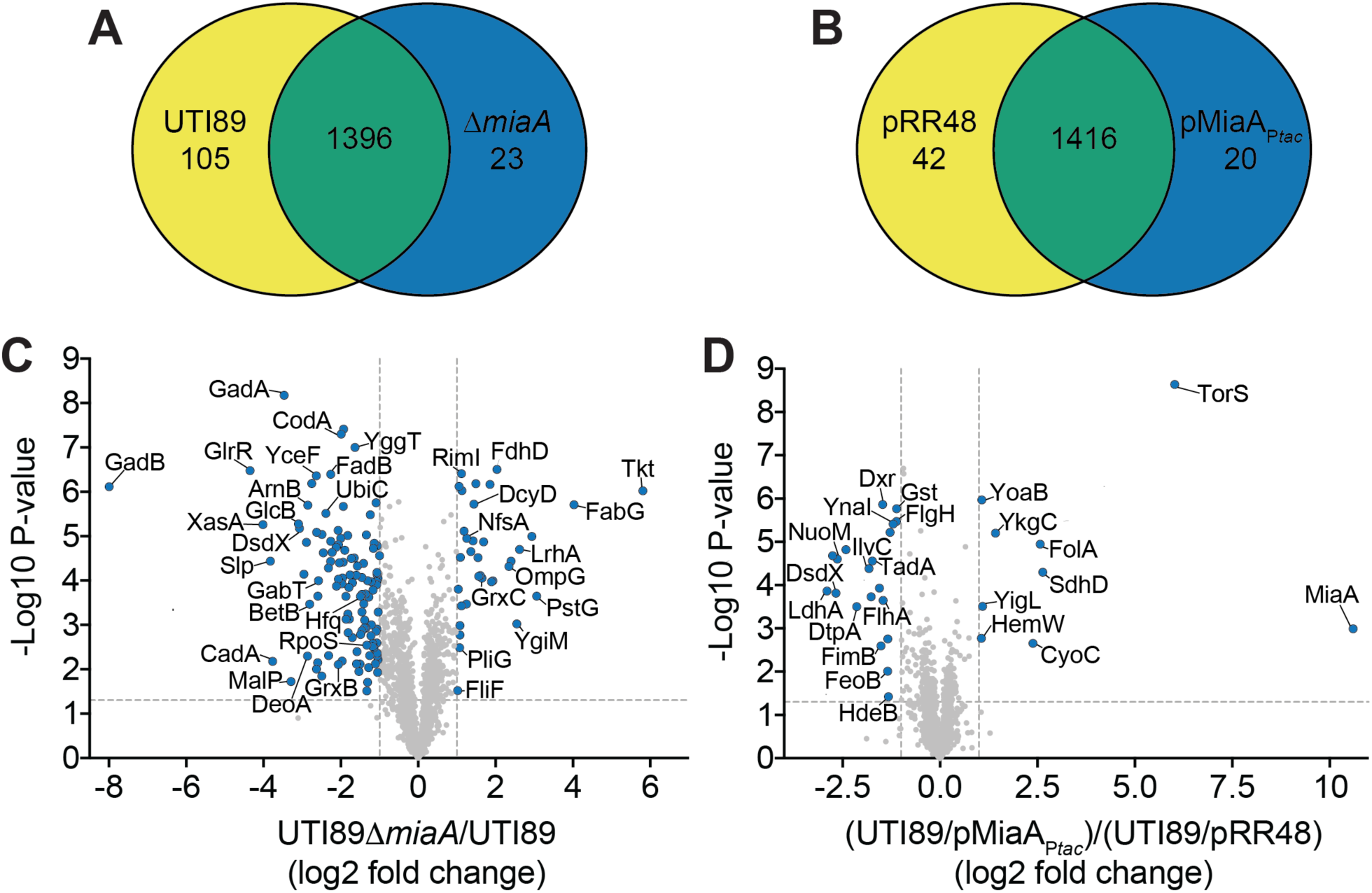
Altering MiaA levels changes the spectrum of expressed proteins. (**A and B**) Venn diagrams indicate numbers of unique and shared proteins detected in wild-type UTI89 versus UTI89Δ*miaA*, or in UTI89/pRR48 versus UTI89/pMiaA_P*tac*_, following growth to OD_600_ of 0.5 in LB. IPTG (1 mM) was included in the UTI89/pRR48 and UTI89/pMiaA_P*tac*_ cultures. Relative protein levels were determined by MudPIT. (**C and D**) Volcano plots show relative protein levels (Log2-fold change) versus P-values (-Log10). Proteins from UTI89Δ*miaA* were quantified relative to the wild-type strain, while proteins levels from UTI89/pMiaA_P*tac*_ were assessed relative to UTI89/pRR48. The vertical dotted lines denote a 2-fold change, while the horizontal dotted lines indicate a *P*-value of 0.05. Blue dots indicate proteins that were significantly changed (*P*<0.05) by at least 2-fold. *P* values were determined by Student’s *t* tests; *n* = 4 independent replicates for each group.

The specific proteins detected, including those that were differentially expressed due to *miaA* deletion or overexpression, are detailed in **Supplemental Dataset S1**. The differentially expressed proteins were assigned to one or more of 14 functional categories (see *Categories* worksheet and embedded graph in **Supplemental Dataset S1**). A majority of the altered proteins were linked with metabolic pathways, secondary metabolites, and functions associated with the bacterial envelope. These included several proteins involved in sugar and fatty acid metabolism and the biosynthesis and regulation of electron transport chains (e.g. UbiC, WrbA, ChrR, Qor, NuoM, NudJ, and CyoC). The dysregulation of these factors likely contributed to the various phenotypic defects observed in our *in vitro* and *in vivo* assays and suggested MiaA involvement in other important processes.

In particular, many of the differentially expressed proteins were shown in previous studies to directly or indirectly affect motility or biofilm development. The former group comprised the chemotaxis protein CheA and the flagella-associated proteins FliF, FlhA, and FlgH. Not unexpectedly, both deletion of the *miaA* gene and MiaA overexpression markedly decreased UTI89 motility on swim plates (Supplemental Fig. S6). MiaB did not affect motility in these assays. Factors linked with biofilm development include YoaB, the type 1 pilus-associated regulator FimB and periplasmic chaperone FimC, the acid stress-response chaperone HdeB, the cellulose synthase catalytic subunit BcsA, and the cytochrome *bo* subunit CyoC. Using yeast extract-casamino acids (YESCA) medium, which promotes the development of elaborate rugose-colony biofilms [77, 78], we found that UTI89Δ*miaA*, but not UTI89Δ*miaB*, formed atypical biofilms with notably less rugosity than the parent strain (Fig. 8). Interestingly, the biofilms formed by UTI89Δ*miaA* were architecturally similar to those formed by a UTI89 mutant lacking the CyoC-interacting partners CyoAB [78].

**Figure 8.**
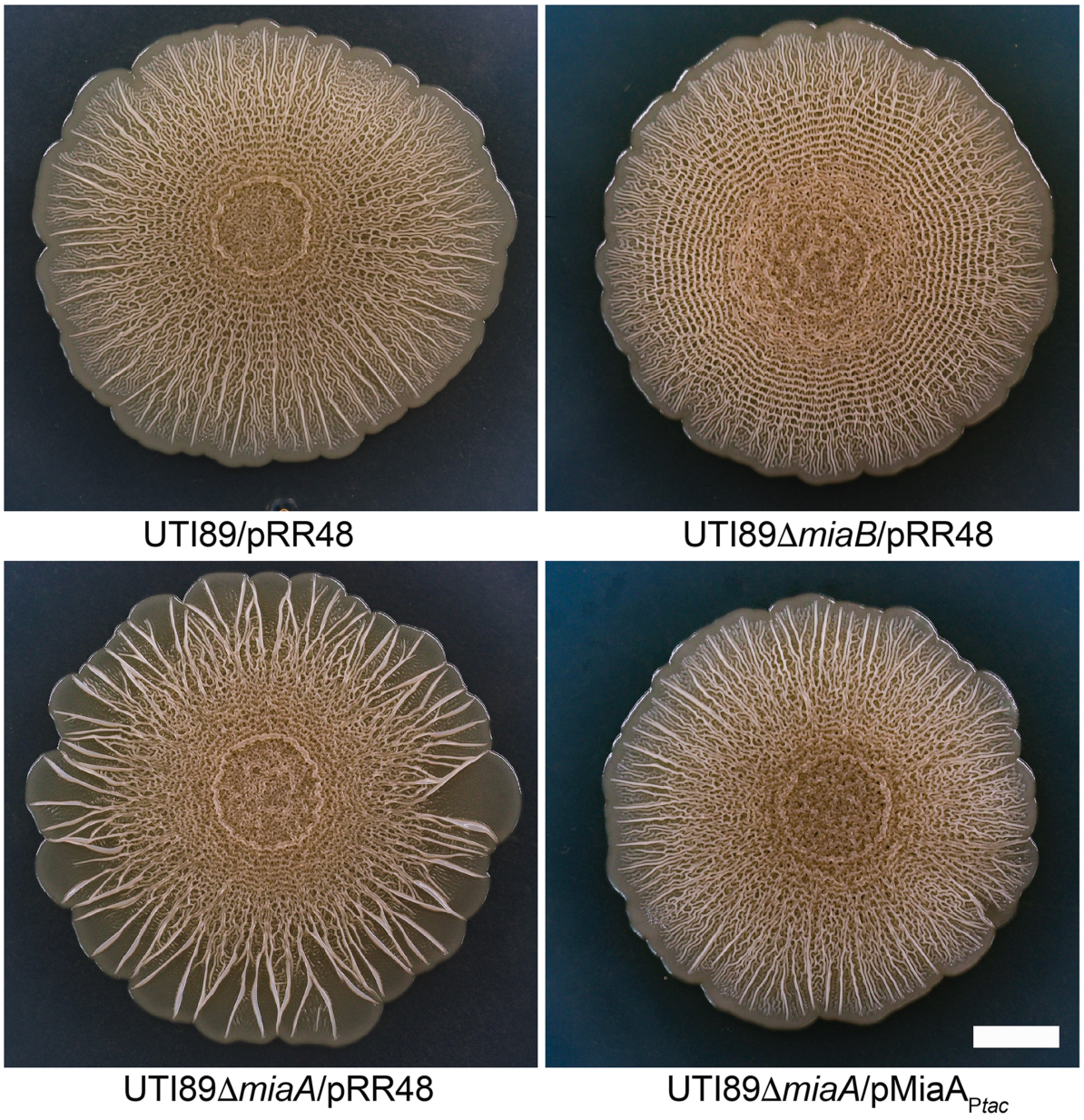
MiaA regulates ExPEC biofilm development. Images show biofilms formed by wild-type UTI89 and its derivatives after 14 days of growth at room temperature on YESCA plates. Photos are representative of at least three independent replicates. Scale bar, 1 cm.

Our MudPIT results also indicated that MiaA can regulate numerous proteins that have been associated with redox and bacterial responses to nitrosative, oxidative, and more generally, genotoxic stresses (**Supplemental Dataset S1**). Aberrant expression of these factors, including proteins like GadB, CadA, Dps, glutathione S-transferase Gst, and the glutatredoxins GrxB and GrxC, may account for increased sensitivity to oxygen and nitrogen radicals (see Fig. 2D-H and Fig. 5D **and** F). Some of these factors, and others like HdeA and HdeB, can also guard against acid stress. Accordingly, follow-up experiments confirmed that UTI89Δ*miaA*, but not UTI89Δ*miaB*, is notably less resistant to acid stress than the wild-type strain (Supplemental Fig. S7). On average, relative to wild-type UTI89, UTI89Δ*miaA* titers were reduced over 6,000-fold following exposure to acidic conditions in LB.

The sensitivity of both UTI89Δ*miaA* and UTI89/pMiaA_P*tac*_ to osmotic stress may arise due to the significant downregulation of proteins like SLP, BetB, YggT, ProP, and YnaI (**Supplemental Dataset S1**). Other differentially expressed proteins that probably contribute to the varied phenotypes associated with *miaA* deletion or MiaA overexpression in our assays include multiple transcriptional regulators, several ribosome- and RNA-associated factors, and the tRNA ligases LysU, TyrS, and PheS. These findings indicate that MiaA is tied into a complex web of factors that can have direct and indirect effects on translation. Driving this point home is the observation that MiaA overexpression suppresses the production of TadA, an enzyme that catalyzes the deamination of adenosine-to-inosine (A-to-I) in Arg2 tRNA and a select set of mRNAs [79, 80]. In K-12 *E. coli,* the A-to-I editing function of TadA can recode at least 12 mRNAs, which results in the generation of proteins with altered activities that can impact bacterial cell physiology [80]. Among the known TadA-edited transcripts is one encoding IlvC, an enzyme involved in isoleucine and valine biosynthesis which, like TadA, is downregulated ∼3.5-fold in UTI89 when MiaA is overexpressed (Fig. 7D).

### UTI89*Δ*miaA phenotypes are not entirely due to aberrant RpoS or Hfq expression

In K-12 *E. coli*, the deletion of *miaA* results in decreased translation of the alternate Sigma factor RpoS (σ^S^) and the small RNA chaperone Hfq [7, 30, 31]. Both of these factors are important for the stress resistance and virulence potential of ExPEC [62, 81]. In line with results from K-12 *E. coli*, our proteomics analysis indicated that RpoS and Hfq levels were reduced 2.5- and 2.8-fold, respectively, in UTI89Δ*miaA* relative to the wild-type strain (Fig. 7C and **Supplemental Dataset S1**). RpoS downregulation in the absence of *miaA* was also confirmed by western blot analysis (Supplemental Fig. S8A) These observations suggest that the phenotypic defects associated with UTI89Δ*miaA* might be attributable to aberrant expression of RpoS or Hfq. However, despite some similarities, the phenotypes that we previously observed with UTI89 mutants lacking either *rpoS* or *hfq* are distinct from one another and from those that we report here with UTI89Δ*miaA* [62, 81]. Furthermore, the induced expression of recombinant RpoS or Hfq (Supplemental Fig. S8B and C) failed to rescue growth of UTI89Δ*miaA* under hyperosmotic conditions (Supplemental Fig. S8D and E). The pRpoS_P*tac*_ and pHfq_P*tac*_ expression constructs used in these assays can complement UTI89 mutants lacking *rpoS* or *hfq*, respectively [62, 81]. Cumulatively, these data indicate that the phenotypes seen with UTI89Δ*miaA* are not entirely due to attenuated expression of either RpoS or Hfq. Also of note, our ability to complement UTI89Δ*miaA* with MiaA expression constructs (see Figs. 2, 3, and 8) demonstrates that the phenotypic defects associated with this knockout mutant are not caused by off target mutations or polar effects on *hfq*, which lies immediately downstream of *miaA*.

### UNN-Leu codon usage by MiaA-sensitive transcripts

Messages like those encoded by *rpoS* and *hfq* are classified as <u>Mo</u>dification <u>T</u>unable <u>T</u>ranscripts (MoTTs), which are identifiable by 1) codon usage different from that of average transcripts and 2) translation that is sensitive to changing levels of tRNA modifications [31, 82]. Studies in the K-12 *E. coli* strain MG1655 of *rpoS*, *hfq*, and other transcripts suggest that MiaA-sensitive MoTTs have higher than average ratios of UNN-Leu codons relative to total Leu codons [30, 31]. This led us to ask if UNN-Leu codon usage correlates with protein expression levels in UTI89 when MiaA is either absent or over-produced. Plotting results from our MudPIT analysis versus UNN-Leu codon usage (**Supplemental Dataset S1**) showed that just over 60% of the proteins that are differentially expressed in UTI89Δ*miaA* or UTI89/pMiaA_P*tac*_ have UNN-Leu codon usage ratios that are greater than the K-12 average ratio of 0.22 (Fig. 9, green dashed line). In line with previous findings [30, 31], RpoS and Hfq are among the differentially expressed proteins with UNN-Leu codon usage ratios of greater than 0.22. However, the average UNN-Leu codon usage ratio in UTI89 is somewhat higher than that in K-12 *E. coli*. Using this value, which is 0.28, less than half of the proteins that are differentially regulated in UTI89Δ*miaA* or UTI89/pMiaA_P*tac*_ have greater than average UNN-Leu codon usage ratios (Fig. 9, black dashed line). Furthermore, among the proteins that are not significantly altered by either deletion or overexpression of *miaA*, about 30% have UNN-Leu codon usage ratios greater than 0.28. Cumulatively, these data indicate that UNN-Leu codon ratios alone may not be especially useful for predicting MiaA-sensitive protein expression patterns within ExPEC strains like UTI89.

**Figure 9.**
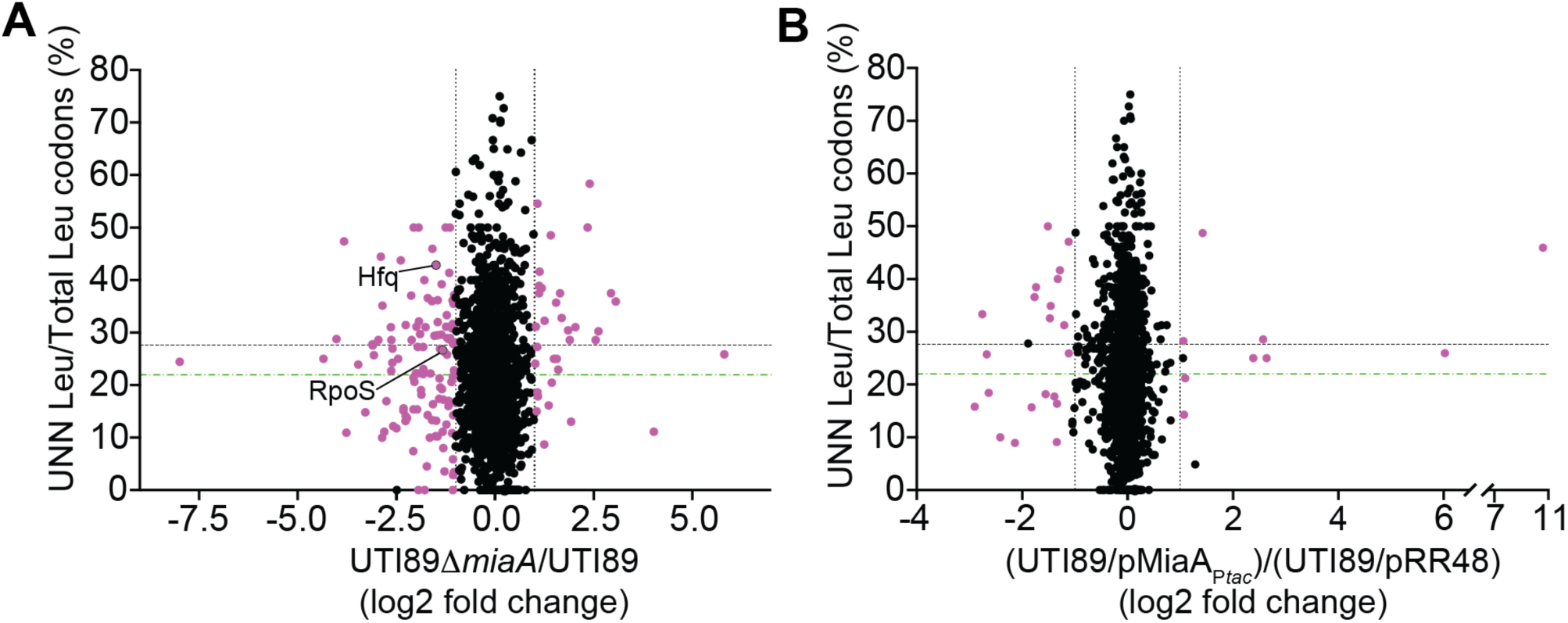
UNN-Leu codon usage ratios vary among MiaA-sensitive transcripts. (**A and B**) Plots show relative protein levels (Log2-fold change) versus UNN-Leu codon usage ratios (UNN Leu/total Leu codons per open reading frame). Proteins in (A) UTI89Δ*miaA* were quantified by MudPIT relative to wild-type UTI89, while proteins levels in (B) UTI89/pMiaA_P*tac*_ were calculated relative to UTI89/pRR48, as in Fig. 7. Purple dots denote proteins that were significantly changed (*P*<0.05, by Student’s *t* tests) by at least 2-fold in UTI89Δ*miaA* or UTI89/pMiaA_P*tac*_ relative to their controls. The vertical dotted lines are placed at the 2-fold change cutoffs. The green and black horizontal dashed lines indicate the average ratios of UNN-Leu codons relative to total Leu codons for all open reading frames encoded by the K-12 strain MG1655 and UTI89, respectively.

## DISCUSSION

The results presented here demonstrate that MiaA is crucial for ExPEC fitness and virulence, and that changing MiaA levels can impact the translation of a broad spectrum of proteins. Our findings are in line with previously published work showing that tRNA modifying enzymes can influence the virulence potential of a variety of microbial pathogens [83]. The attenuation of bacterial virulence-related phenotypes in the absence of a specific tRNA modifying enzyme can, in some cases, be explained by sub-optimal translation of specific toxins or key regulatory factors [14, 19–21]. For example, deletion of *miaA* in the diarrheagenic bacteria *Shigella flexneri* ablates translation of the transcriptional master regulator VirF, resulting in the reduced expression of downstream virulence factors [11, 84]. Overexpression of recombinant VirF alone is sufficient to rescue the *miaA* mutant, suggesting that low-level production of VirF is in large part responsible for the virulence-related defects caused by the deletion of *miaA* in *S. flexneri*. In contrast, our work indicates that the diverse phenotypes affected by MiaA expression in the ExPEC isolate UTI89 are not attributable to any single factor, but rather arise due to the altered expression and dysregulation of multiple proteins and pathways downstream of MiaA.

The ms^2^i^6^A modification is understood to affect the fidelity of translation [5, 8, 27]. Earlier work in K-12 *E. coli* and *Salmonella* strains showed that bacteria lacking *miaA* have an increase in the +1 direction of frameshifting, but not the −1 direction [73, 74]. In these studies, the i^6^A modification was found to be a major contributor to ribosome fidelity. In UTI89, significant increases in frameshifting were seen in both the +1 and −1 directions when *miaA* was knocked out. When MiaA was overproduced, we also observed marked elevation of frameshifting in the −1 direction, while frameshifting levels in the +1 direction were more modest. Most reports to date indicate that tRNA modifications typically affect frameshifting primarily in one direction [5, 74]. At first glance, our data seemingly counter this trend. However, we note that the expression of firefly luciferase downstream of the HIV-derived linker in our reporter system may also occur as a consequence of a +2 frameshift, rather than a −1 frameshift, which would more closely mirror what was observed in previous studies with K-12 *E. coli* and *Salmonella* strains [73, 74]. It is also possible that the apparent increases in both −1 and +1 frameshifting observed in our assays reflect the presence of MiaA-sensitive regulatory circuits in UTI89 that are different from those in K-12 *E. coli* or *Salmonella* strains.

During the course of this study, we were surprised to observe that MiaA levels in the ExPEC reference strain UTI89 were substantially decreased in response to high salt (see Fig. 4). Ongoing work indicates that MiaA levels in UTI89 are also altered upon exposure to other stressors, such as MV (Supplemental Fig. S9A and B). Due to the sweeping phenotypes seen in the absence of MiaA, we hypothesized that the levels of MiaA would have stayed the same or increased in response to stressors like high salt and MV. However, our data indicate that MiaA levels are fine-tuned within ExPEC such that too much or too little enzyme can have similarly detrimental consequences. The effects of MiaA expression in our growth curve assays were dose–dependent, with high-level expression of MiaA being nearly as disruptive as the deletion of *miaA*. For instance, low-level expression of MiaA restored the resistance of UTI89Δ*miaA* to high salt, whereas overexpression of MiaA resulted in greatly increased sensitivity (see Fig. 5).

MiaA is part of a complex superoperon and its regulation, and the regulation of tRNA modifying enzymes in general, is not well understood [33, 85, 86]. Though we did not investigate MiaA regulation in detail here, our RT-qPCR experiments indicate that MiaA levels are reduced in response to high salt stress via a post-transcriptional mechanism (Fig. 4C). Interestingly, *miaA* has a higher-than-average UNN Leu codon usage ratio of 0.46, suggesting that MiaA may help regulate the translation of its own transcripts [31]. Furthermore, we note that MiaA levels are intensified in the presence of the metal chelator EDTA (Supplemental Fig. S9C), raising the possibility that the quantities of this tRNA modifying enzymes are controlled by one or more EDTA-sensitive metalloproteases. The factors that modulate MiaA levels during times of stress require further investigation.

By adjusting the levels of tRNA modifying enzymes like MiaA, ExPEC and other organisms may be able to vary the diversity of translated proteins and thereby optimize adaptive responses to stressful stimuli [14, 33, 46, 87–90]. Indeed, the overexpression and deletion of *miaA* led to the generation of distinct proteomes by UTI89 (Fig. 7) and compromised the ability of this ExPEC strain to deal with multiple stressors. Because tRNA modifications can have pleiotropic effects, it is not always easy to distinguish the direct and indirect effects that tRNA modifying enzymes like MiaA have on translation [90]. For example, pioneering work in K-12 *E. coli* indicates that the efficient translation of RpoS and Hfq relies on MiaA for proper decoding of UNN-Leu codons [7, 30, 31], but these factors can themselves regulate the expression of numerous other proteins [91–93]. The capacity for MiaA to have additional, indirect effects on the fidelity and specificity of translation is further highlighted by our proteomics data showing that MiaA impacts the expression of multiple ribosome- and RNA-associated factors, tRNA ligases, and the RNA editing enzyme YfhC (TadA). These findings suggest the existence of a complex network of RNA and translational modifiers that can regulate the expression of one another. Layered on top of this is the potential for MiaA to affect the biosynthesis and availability of specific metabolites used by other tRNA modifying enzymes [15, 33].

Increases in frameshifting due to changing levels of MiaA may also allow for more error-prone translation and the subsequent diversification of expressed proteins, which could allow bacteria to better deal with stressful stimuli. The ability of cells to actively regulate frameshifting and other translational errors in order to generate mutant proteins that deviate from those encoded by the genome is gaining appreciation as an adaptive response to stress [94–97]. Ongoing studies aim to utilize ribosomal profiling along with RNA-seq and proteomics to determine if off-frame and mutant proteins are being produced by ExPEC via translational modifiers like MiaA in response to stressful stimuli. Similar lines of research may also shed light on the somewhat more cryptic functions of the MiaB-catalyzed tRNA modification.

In the absence of *miaA*, the i^6^A and ms^2^i^6^A modifications are not detectable (Fig. 4), as expected from previously published work [86]. When MiaA is overproduced, high levels of i^6^A are observed, while ms^2^i^6^A modification levels remain relatively stable (Fig. 5A). This suggests that only a fraction of the UNN-decoding tRNAs are fully modified in the cell at any time. Interestingly, high salt stress reduces both MiaA expression and ms^2^i^6^A levels, but does not significantly affect i^6^A levels (Fig. 4). In contrast, the disruption of *miaB* prevents the formation of ms^2^i^6^A and causes marked increases in i^6^A levels (Supplemental Fig. S4), but this had no phenotypic effect in any of our assays. These findings present a conundrum – why does high-level production of i^6^A due to MiaA overexpression attenuate the stress resistance of UTI89 while even higher levels of i^6^A that build up in the absence of MiaB had no overt phenotypic effects in our assays? In considering this issue, it should be noted that we quantified relative levels of i^6^A and ms^2^i^6^A, and not specific tRNAs, leaving open the possibility that changing levels of MiaA differentially affect distinct tRNA subsets. This alone could account for the contrasting phenotypic effects linked with elevated i^6^A levels due to MiaA overexpression versus those caused by *miaB* deletion. MiaA overexpression may also be detrimental due to depletion of substrates like dimethylallyl diphosphate (DMAPP) that feed into other critical pathways, including the biosynthesis of ubiquinone. The deletion of *miaB*, by causing a buildup of i^6^A rather than accelerated production of this modified residue, may have less of an abrupt impact on the availability substrates like DMAPP. Finally, it is feasible that MiaA has moonlighting function(s), affecting non-tRNA targets and compromising bacterial fitness when produced in excess. Cumulatively, the findings presented here highlight the central and complex roles that core metabolic genes like *miaA* can have on the elaboration and fine-tuning of pathogen stress resistance and virulence-associated phenotypes.

## EXPERIMENTAL PROCEDURES

### Bacterial strains

Strains used in this study are listed in Supplemental Table S1. Mutant strains were constructed in the reference ExPEC isolate UTI89 using the lambda Red recombination system and primers detailed in Supplemental Table S2, as previously described [48]. The chloramphenicol resistance (Clm^R^) cassette flanked by LoxP sites was amplified from plasmid pKD3 using primers that contain overhanging ends with ∼40 bp of homology near the 5’ and 3’ ends of each target locus. PCR products were introduced by electroporation into UTI89 carrying pKM208, which encodes an IPTG-inducible lambda Red recombinase [49]. Knockouts were verified by PCR using primers indicated in Supplemental Table S2.

### Plasmids

Expression and reporter constructs were generated using standard molecular biology approaches and primers listed in Supplemental Table S2. The *miaA* and *miaB* genes were amplified from UTI89 by PCR, digested, and ligated into pRR48 using Pst1 and Kpn1 restriction sites to create pMiaA_P*tac*_ and pMiaB_P*tac*_. Sequences encoding Hfq fused with C-terminal 6xHis and Flag tags were cloned using a similar approach to make pHfq_P*tac*_. To create pMiaA_nat_, the UTI89 *miaA* sequence was amplified along with 200 base pairs of flanking sequences, including the *miaA* promoter, and then ligated into the EcoR1 site of pACYC184. The plasmid pMiaA-Flag_nat_, having the *miaA* promoter region upstream of sequences encoding MiaA with a C-terminal Flag-tag, was produced similarly.

The dual-luciferase reporter plasmids used for the frameshifting assays were created using p2Luc plasmids as templates [75, 76]. The genes encoding the renilla and firefly luciferases were amplified by PCR along with intergenic Az1- or HIV-derived linker sequences. A Shine-Dalgarno ribosome binding site was incorporated into the forward primer (p2Luc_F) primer to promote translation of the linked luciferases. PCR products were digested and ligated into the KpnI and HindIII sites of pBAD18 (Ap^R^) and pBAD33 (Cam^R^). Plasmids with different resistance cassettes were needed for use with UTI89Δ*miaA* (Cam^R^) and UTI89 carrying pMiaA_P*tac*_ or the empty vector pRR48. The Az1- and HIV-derived linker region sequences are noted in Fig. 6A, and were chosen because they contain MiaA-sensitive UNN codons. Control plasmids in which the Az1 and HIV linkers are altered to place the two luciferases in-frame were generated in an analogous fashion using previously described p2Luc plasmids as templates [75, 76].

### Bacterial growth analysis

UTI89 and its derivatives were grown from frozen stocks in 5 ml of LB, 100 mM MES-buffered LB (MES-LB; pH 5.0), or modified M9 medium (6 g/liter Na_2_HPO_4_, 3 g/l KH_2_PO_4_, 1 g/l NH_4_Cl, 0.5 g/l NaCl, 1 mM MgSO_4_, 0.1 mM CaCl_2_, 0.1% glucose, 0.0025% nicotinic acid, 0.2% casein amino acids, and 16.5 µg/ml thiamine in H_2_O) at 37°C overnight in loosely capped 20-by-150-mm borosilicate glass tubes with shaking (225 rpm, with tubes tilted at a 30° angle). Overnight cultures were brought to an OD_600_ of ∼1.0 and then sub-cultured 1:100 into LB, MES-LB, or M9 medium. Growth curves were acquired using a Bioscreen C instrument (Growth Curves USA) with 200-μl cultures in 100-well honeycomb plates shaking at 37°C. Cultures included extra NaCl (5% w/v), 1 mM MV (Sigma-Aldrich), 1 or 2 mM ASN (Sigma-Aldrich), or IPTG, as indicated. MV and ASN solutions were prepared fresh just before use. All growth curves were determined using quadruplicate samples with at least three independent replicates. Overnight cultures of strains carrying plasmids for complementation experiments were grown in the presence of antibiotics (100 μg of ampicillin/ml or 50 μg of tetracycline/ml) to maintain the plasmids, but antibiotics were not included in media used for the subsequent growth assays.

### Mouse models

All animals used in this study were handled in accordance with protocols approved by the Institutional Animal Care and Use Committee at the University of Utah (Protocol number 10-02014), following US federal guidelines indicated by the Office of Laboratory Animal Welfare (OLAW) and described in the Guide for the Care and Use of Laboratory Animals, 8th Edition. Mice were purchased from The Jackson Laboratory, housed 3 to 5 per cage, and allowed to eat (irradiated Teklad Global Soy Protein-Free Extruded chow) and drink antibiotic-free water *ad libitum*.

### Competitive gut colonization assays

For these assays, a kanamycin resistance cassette (Kan^R^) was inserted into the *att*Tn7 site of UTI89 to create UTI89::Kan^R^, which can be easily distinguished from the chloramphenicol resistant (Cam^R^) *miaA* and *miaB* knockout mutants by plating on selective media. Previous work demonstrated that insertion of resistance cassettes into the *att*Tn7 site does not impact ExPEC fitness within the gut [55, 56]. Individual cultures of UTI89::Kan^R^ (standing in as the wild-type strain), UTI89Δ*miaA*, and UTI89Δ*miaB* were grown statically from frozen stocks for 24 h at 37°C in 250-ml flasks containing 20 ml of modified M9 medium. Each knockout mutant was then mixed 1:1 with UTI89::Kan^R^ (6 ml of each culture) and then pelleted by centrifugation at 8,000 x *g* for 8 minutes at room temperature. The bacterial pellets were then washed once with phosphate-buffered saline (PBS), pelleted again, and resuspended in 0.5 ml of PBS. Female SPF BALB/c mice aged 7 to 8 weeks were inoculated via oral gavage with 50 μl PBS containing ∼10^9^ CFU of each bacterial mixture. At the indicated time points post-inoculation, individual mice were placed into unused takeout boxes for a few minutes for weighing and feces collection. Freshly deposited feces were collected from the boxes and immediately added to 1 ml of 0.7% NaCl, weighed, and set on ice. The samples were then homogenized and briefly centrifuged at low speed to pellet any insoluble debris. Supernatants were serially diluted and plated onto LB agar containing either chloramphenicol (20 μg/ml) or kanamycin (50 μg/ml) for selective growth of UTI89::Kan^R^ (wild type), UTI89Δ*miaA*, or UTI89Δ*miaB*. Fecal samples were also analyzed prior to the start of each experiment to ensure that there were no endogenous bacteria present that were resistant to chloramphenicol or kanamycin. CIs were calculated as the ratio of knockout over wild-type bacteria recovered in the feces divided by the ratio of knockout over wild-type bacteria present in the inoculum [55, 57]. A total of 7 to 8 mice in two independent assays were used for each set of bacterial strains tested.

### UTI model

The murine UTI model was used essentially as described by our group and others [98, 99]. Wild-type UTI89 and the *miaA*, and *miaB* knockout mutants were grown from frozen stocks in 20 ml LB broth in 250 mL Erlenmeyer flasks without shaking at 37°C for 24 hours. Bacteria were then pelleted by centrifugation (8 minutes at 8,000 x g) and resuspended in PBS. Seven- to eight-week-old female CBA/J or C3H/HeJ mice were briefly anesthetized by isoflurane inhalation and slowly inoculated via transurethral catheterization with 50 µL of PBS containing a suspension of ∼10^7^ bacteria. Bacterial reflux into the kidneys using this procedure is rare, occurring in less than 1% of the test animals. At 0.25, 1, 3, or 9 days post-inoculation, mice were sacrificed and bladders were harvested aseptically, weighed, and homogenized in 1 ml PBS containing 0.025% Triton X-100. Bacterial titers within the homogenates were determined by plating serial dilutions on LB agar plates. Nine or more mice in total, from two independent experiments, were used for each bacterial strain and time point examined.

### Sepsis model

UTI89, UTI89Δ*miaA*, and UTI89Δ*miaB* were grown from frozen stocks in 20 ml M9 broth without shaking at 37°C for 24 h, pelleted by centrifugation at 8,000 x *g* for 8 minutes, and washed once with PBS, pelleted again, and resuspended in PBS. Seven- to eight-week-old female C57Bl/6 mice were briefly anesthetized by isoflurane inhalation and infected via intraperitoneal injection of ∼10^7^ CFU within 200 μl PBS. Mice were monitored over a 72-hour period for signs of morbidity and mortality. Alternatively, at 6 hours post-inoculation mice were sacrificed and the liver, kidneys, and spleens were harvested aseptically, weighed, and homogenized in 1 ml PBS containing 0.025% Triton X-100. Bacterial titers within the homogenates were determined by plating serial dilutions on LB agar plates.

### Invasion, adhesion, and intracellular persistence assays

Bacteria were grown at 37°C for 48 h in 20 mL static LB broth to induce expression of type 1 pili, which are important mediators of UPEC adherence and entry into host cells [99]. Host cell association and gentamicin protection-based invasion and overnight intracellular persistence assays were performed as previously described using the human bladder epithelial cell line 5637 (HTB-9; ATCC) [100]. Of note, UTI89Δ*miaA* is about 3-fold more sensitive to the host-cell impermeable antibiotic gentamicin, as determined by using Etest Strips (VWR) (Supplemental Fig. S10). However, this likely had no effect on results from the cell culture-based invasion and intracellular survival experiments, as the concentrations of gentamicin (100 and 10 μg/ml) used in these assays exceed those needed to effectively kill extracellular UTI89, UTI89Δ*miaB*, and UTI89Δ*miaA*.

### Biofilm analysis

*In vitro* rugose biofilm assays were performed starting with cultures grown overnight at 37°C shaking in LB, as described [78]. Bacteria from each culture were then brought to an OD_600_ of ∼1.0 and 10 μl aliquots were spotted onto YESCA agar plates (12 g/l Casamino acids, 1.2 g yeast extract, 22 g agar) and incubated at RT (∼20-22°C). After 14 days, biofilm images were acquired by focus stacking using an M.Zuiko Digital ED 60 mm lens mounted on an Olympus OM-D E-M1 Mark II camera.

### Motility assays

Cultures of UTI89, UTI89Δ*miaA*, and UTI89Δ*miaB* grown overnight shaking in LB or M9 medium were brought to OD_600_ of 1.0. Swim motility plates, containing 0.2% agar in LB or M9 medium, were inoculated with 2 µl of each bacterial suspension delivered just below the agar surface. The diameter of bacterial spreading was measured every 1-2 hours over the course of an 8-10 hour-incubation at 37°C. Swim rates were calculated during logarithmic growth. To assess the effects of MiaA and MiaB overexpression on motility, tryptone soft agar plates [101] containing 50 μg/ml ampicillin and 100 μM IPTG were inoculated with UTI89/pRR48, UTI89/pMiaA_P*tac*_, and UTI89/pMiaB_P*tac*_ from overnight shaking cultures. Plates were imaged after a 6-hour incubation at 37°C.

### Acid resistance assays

Bacterial strains from overnight cultures were diluted 1:100 in fresh LB and grown shaking at 37°C for 3 h. Concentrated HCl was then added to each culture to adjust the pH to 3.0 and incubations were continued for another 30 minutes. Bacteria from 1 ml of each culture were then pelleted at 16,000 x *g* for 5 min and washed in PBS. Surviving bacteria were enumerated by plating serial dilutions on LB agar and normalized to input titers.

### Osmotic stress resistance assays

UTI89/pACYC184, UTI89Δ*miaA*/pACYC184, UTI89Δ*miaA*/pMiaA_nat_, and UTI89Δ*miaB* were grown shaking overnight at 37°C in 5 ml LB broth with 20 µg/ml tetracycline and then back diluted 1:100 into 5 ml fresh LB (+ tetracycline). After 5 h shaking at 37°C, a 1-ml aliquot of each culture was pelleted, resuspended in 1 ml of sterile water with or without 0.1% glucose, and incubations were continued for another 2 h with shaking at 37°C. Viable bacteria present at 0, 30, 60, 90, and 120 min after resuspension in water were quantified by dilution plating and normalized to input titers. Growth curves in LB ± 5% NaCl were acquired as described above.

### Western blot analysis

Bacterial pellets were frozen at −80°C and then resuspended in B-PER lysis reagent (Thermo Scientific) supplemented with 1 mM phenylmethylsulfonyl fluoride, protease inhibitor cocktail (Roche), and Lysonase Bioprocessing Reagent (Novagen). After a 15-minute incubation at room temperature, samples were spun for 1 minute at 13,000 x *g* to remove large cell debris, and protein concentrations in the supernatants were determined using the BCA reagent system (Pierce). Equivalent protein amounts were resolved by SDS-PAGE and subsequently transferred to Immobilon PVDF-FL membranes (Millipore). Blots were probed using mouse anti-Flag M2 (1:3000; Sigma-Aldrich), rabbit anti-Flag (Immunology Consultants laboratory, inc.), and mouse anti-RpoS (anti-SigmaS; Biolegend) and visualized using enhanced chemiluminescence with HRP-conjugated secondary antibodies (1:3000 or 1:5000; Amersham Biosciences), as described [102]. To ensure that equivalent amounts of protein from each sample were analyzed, blots were re-probed using rabbit anti-*E. coli* antisera (1:2,000 or 1:5000; BioDesign International).

### Analysis of relative i6A and ms2i6A levels

UTI89 and UTI89Δ*miaA* were grown from frozen stocks shaking at 37°C overnight in LB. UTI89/pRR48 and UTI89/pMiaA_P*tac*_ were grown similarly using LB supplemented with ampicillin (100 μg/ml). The bacteria were sub-cultured 1:100 into 6 ml of LB ± 1 mM IPTG and then grown shaking to an OD_600_ of 0.5. After adjusting the cultures to OD_600_ of 1.0, the bacteria were pelleted by spinning at 8000 x *g* for 1.5 minutes. Pellets were then resuspended in 1 ml of RNA*later* Stabilization Solution (ThermoFisher) and stored at 4°C overnight prior to extraction of RNA using a Norgen Total RNA Extraction Kit.

Samples were analyzed using a Hypersil GOLD C18 column (2.1 mm × 150 mm, 1.9 μm particle size; Thermo Fisher) attached to a Thermo Scientific Dionex UltiMate 3000 UHPLC instrument in line with an LTQ-OrbiTrap XL instrument (Thermo Fisher). The LC-MS parameters were based upon a procedure described previously [103, 104], with the following adjustments. The UHPLC column was pre-equilibrated in 100% Buffer A [50 mM ammonium acetate (Fisher) in LC–MS Optima water]. Buffer B consisted of 60% (v/v) LC–MS Optima acetonitrile (Fisher) and 40% LC–MS water (Fisher). The reaction components were eluted at a rate of 0.2 ml/minute with the following program: 0% B from 0 to 3.46 min, 0 to 0.9% B from 3.46 to 3.69 min, 0.9 to 1.5% B from 3.69 to 3.92 min, 1.5 to 3% B from 3.92 to 4.25 min, 3 to 20% B from 4.25 to 6.5 min, 20 to 25% B from 6.5 to 7 min, 25 to 40% B from 7 to 8.5 min, 40 to 45% B from 8.5 to 9.25 min, 45 to 60% B from 9.25 to 9.95 min, 60 to 100% B from 9.95 to 10.45 min, 100% B from 10.45 to 16 min, 100 to 0% B from 16 to 16.1 min, and 0% B from 16.1 to 20 min. The flow from the column was diverted to the mass spectrometer from 3.5 minutes to 17 minutes during the UHPLC program. The mass spectrometer was operated in positive ion mode, and authentic guanosine material (Sigma Aldrich) was used to generate a tune file for the instrument. The observed *m/z* values of the +1 charge states of the i6a and ms2i6a RNA bases were 336.1658 and 382.1535, respectively. The observed retention times for i6A and ms2i6a were determined from the center of their extracted ion chromatogram peaks to be 15.55 and 16.45 minutes, respectively. The retention time of i6a from biological extracts was consistent with the retention time of authentic i6a material (Cayman Chemical). Absolute intensities of the i6a ions were retrieved from the mass spectrum scanned between 15.4 – 16.1 minutes, while the absolute intensities of the ms2i6a ions were retrieved from the mass spectrum scanned between 16.2 and 17.0 minutes. The scan range was chosen to include the entire peak of an EIC trace, excluding mass spectral data recorded out of these bounds. This ensured that the intensities of ions 336.17 and 382.15 arise from eluted i^6^A and ms^2^i^6^A material and did not include background during the rest of the run. The scan windows were wide enough to account for any small drift in retention that might occur from sample to sample. Because total RNA concentration varied by sample, the samples were normalized against the total RNA concentration of each sample, as estimated via a NanoDrop measurements at 260 nm.

### Quantification of frameshifting

UTI89, UTI89Δ*miaA,* UTI89/pRR48 and UTI89/pMiaA_P*tac*_ carrying one of the dual-luciferase reporter plasmids (see Supplemental Table S1) were grown overnight in LB supplemented with chloramphenicol (20 μg/ml) or ampicillin (100 μg/ml). The cells were sub-cultured 1:100 into 6 ml LB with and without 1mM IPTG. At an OD_600_ of 0.2, arabinose (0.2%) was added to all of the cultures to induce expression of the luciferases. Cells were allowed to continue growing until reaching an OD_600_ of 0.5, at which point the cultures were adjusted to an OD_600_ of 1.0 and pelleted by spinning at 8000 x *g* for 1.5 minutes. The pellets were then subjected to one freeze-thaw cycle before being resuspended in Passive Lysis Buffer (Promega; E1910). One scoop of 0.15 mm zirconium oxide beads (Next Advance; ZrOB015) was added to each tube and bacteria were lysed using a Bullet Blender (Next Advance) set at speed 8 for 3 min. After a 30-second spin in a microfuge to pellet beads and any large debris, the supernatants were collected and luciferase activities were analyzed as previously described [76]. Briefly, the Dual-Luciferase Reporter Assay System (Promega) was used in combination with the Veritas Microplate Luminometer from Turner Biosystems to quantify activity of the two luciferases. Frameshifting was calculated by first determining the ratio of firefly to renilla luciferase activity for each sample, and then normalizing each out-of-frame construct (pCWR43 and pCWR45) with their associated in-frame control (pCWR42 and pCWR44, respectively).

### Proteomics

UTI89, UTI89Δ*miaA,* UTI89/pRR48, and UTI89/pMiaA_P*tac*_ were grown to mid-log phase (OD_600_∼0.5) in LB shaking at 37°C. IPTG (1 mM) was included for UTI89/pRR48 and UTI89/pMiaA_P*tac*_. About 1X10^9^ CFU from each culture was pelleted at 8,000 x *g* for 1.5 minutes. Supernatants were then removed and cells were plunged into liquid nitrogen and subsequently analyzed using MudPIT with the MSRC Proteomics Core at Vanderbilt University. Label-free quantification (LFQ) values were loaded into Prostar software for statistical analysis and visualization. The data set was filtered by requiring all conditions to contain at least two values. Imputation for partially observed values was done with the Structured Least Square Adaptative algorithm. Imputation for conditions in which values were missing for a specific protein in all three biological replicates used the DetQuantile algorithm with the settings Quantile:2.5 and Factor:1. Statistical analysis was performed using the 1vs1 settings and Student’s *t*-tests. Differentially expressed proteins were categorized (**Supplemental Dataset S1**) based on literature searches and information drawn from EcoCyc ([105]; http://ecocyc.org/), STRING Protein-Protein Interaction Networks Functional Enrichment Analysis ([106]; https://string-db.org/), and Phyre2 ([107]; http://www.sbg.bio.ic.ac.uk/~phyre2/html/page.cgi?id=index). The proteomics output files will be uploaded to ProteomeXchange.

### RT-qPCR analysis

UTI89 was diluted 1:100 from overnight cultures into fresh LB, grown shaking for 2.5 hours at 37°C prior to resuspension in LB or LB + 5% NaCl. After another one-hour incubation bacteria were pelleted and total RNA was extracted using the miRNeasy mini kit (QIAGEN). RNA samples were treated with RNase-Free DNase (QIAGEN) and cDNA was made using SuperScript IV VILO Master Mix (Invitrogen) according to the manufacturer’s protocol. Quantitative PCR (qPCR) was carried out using primers listed in Supplemental Table S2 with the PowerUp SYBR Green Master Mix (Thermo Fisher Scientific) on a QuantStudio 3 Real-Time PCR Instrument (Applied Biosystems). Replicas were made for each cDNA sample and *miaA* and *miaB* levels were normalized to *rpoD*. Products were resolved in 1.5% agarose gels, stained with ethidium bromide, and visualized using a GelDoc system (BioRad Technologies) to help verify the specificity of the RT-qPCR results.

### Codon usage analysis

Codon frequencies (UNN Stats tab, **Supplemental Dataset S1**) were calculated for each gene in UTI89 (accessions CP000243.1 and CP000244.1) using custom Python scripts that leverage the BioPython and NumPy packages.

### Statistical analysis

*P* values were determined as indicated by Log-Rank (Mantel-Cox), Mann-Whitney U tests, ANOVA, or Student’s *t*-tests performed using Prism 9.0.0 software, with corrections as indicated (GraphPad Software). Data distribution normality (Gaussian) was not assumed, such that non-parametric tests were used for the mouse experiments. *P*-values of less than or equal to 0.05 were defined as significant.

## ACKNOWLEDGEMENTS

We thank W. Hayes McDonald of the Vanderbilt School of Medicine Mass Spectrometry Research Center for help with the proteomics. This study was funded in part by NIH grants to M.A.M. (GM134331, AI135918, AI095647, and AI088086) and to V.B. (GM126956). M.G.B. was supported by T32 AI055434 from the National Institute of Allergy and Infectious Diseases, and A.J.L. was supported by T32 DK007115 from the National Institute of Diabetes and Digestive and Kidney Diseases. The authors have no conflicts of interest to declare.

## Author Contributions

MGB, BAF, and MAM designed, supervised the study, performed research, analyzed the data, and drafted the paper. WMK, AJL, JRB, MH, VB, and MTH helped with design, experimentation, and editing. AT, CJB, LMB, and QZ performed experiments and helped process samples. All authors contributed to and approved the submission of this paper.

## SUPPLEMENTAL INFORMATION

**Supplemental Figure S1:**
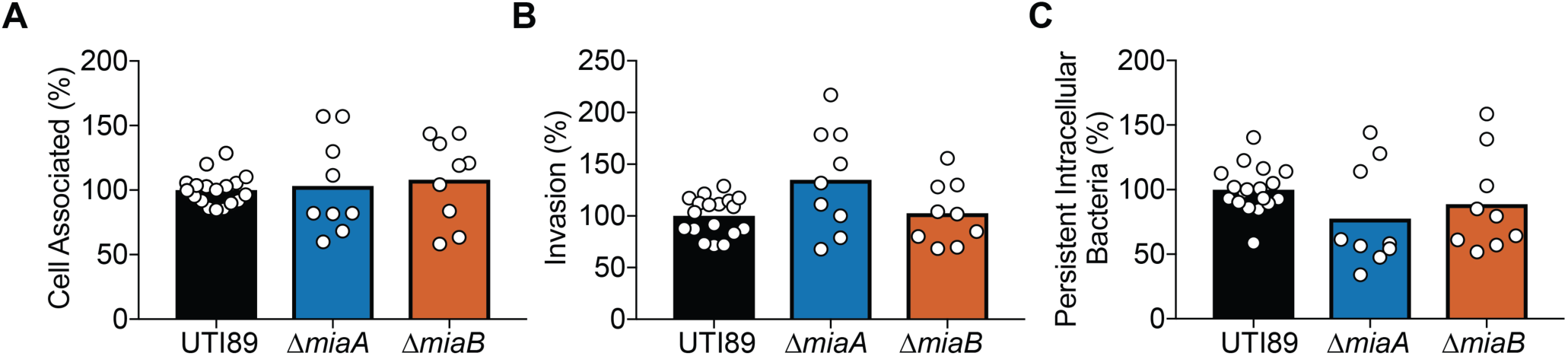
MiaA and MiaB do not affect the ability of ExPEC to bind, invade, or persist intracellularly within bladder epithelial cells. Human bladder epithelial cells (5637 cells) were infected with UTI89, UTI89Δ*miaA*, or UTI89Δ*miaB* for 2 h, followed by a second 2-h incubation in the presence of the bactericidal, host cell-impermeable antibiotic gentamicin (100 μg/ml). Graphs show (**A**) the levels of host cell-associated bacteria prior to the addition of gentamicin, (**B**) and the relative numbers of intracellular bacteria recovered after the 2-h incubation in media containing gentamicin. (**C**) Longer-term bacterial persistence within the bladder cells was assessed by continued incubation of infected host cells for an additional 12 h with gentamicin. For the longer persistence assays, a submaximal concentration of gentamicin (10 μg/ml) was used to prevent extracellular growth of UPEC while limiting possible leaching of the antibiotic into the host cells. Data are expressed relative to wild-type UTI89, with bars indicating median values from 9 to 18 independent experiments performed in triplicate.

**Supplemental Figure S2.**
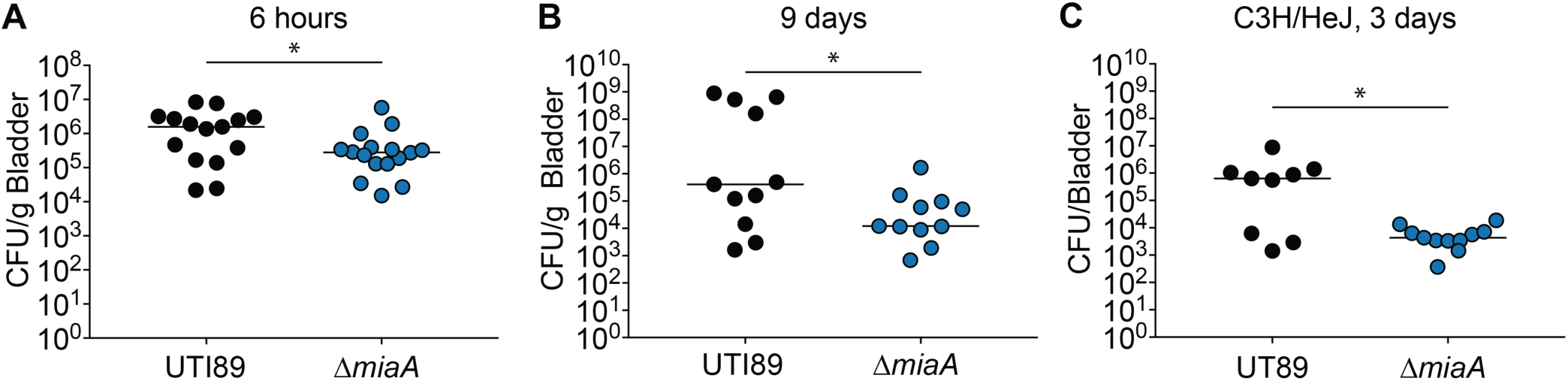
MiaA promotes ExPEC colonization and persistence within the murine bladder. (**A and B**) The bladders of adult female CBA/J mice were inoculated via transurethral injections with ∼10^7^ CFU of wild-type UTI89 or UTI89Δ*miaA*. Mice were sacrificed (A) 6 hours or (B) 9 days later and bacterial titers within the bladders were determined by plating tissue homogenates. (**C**) Graph shows bacterial titers recovered from the bladders of adult female C3H/HeJ mice 3 days after inoculation with UTI89 or UTI89Δ*miaA*. Bars in all graphs denote median values. *, *P <* 0.05 by Mann Whitney U tests; *n* ≥ 9 mice per group.

**Supplemental Figure S3.**
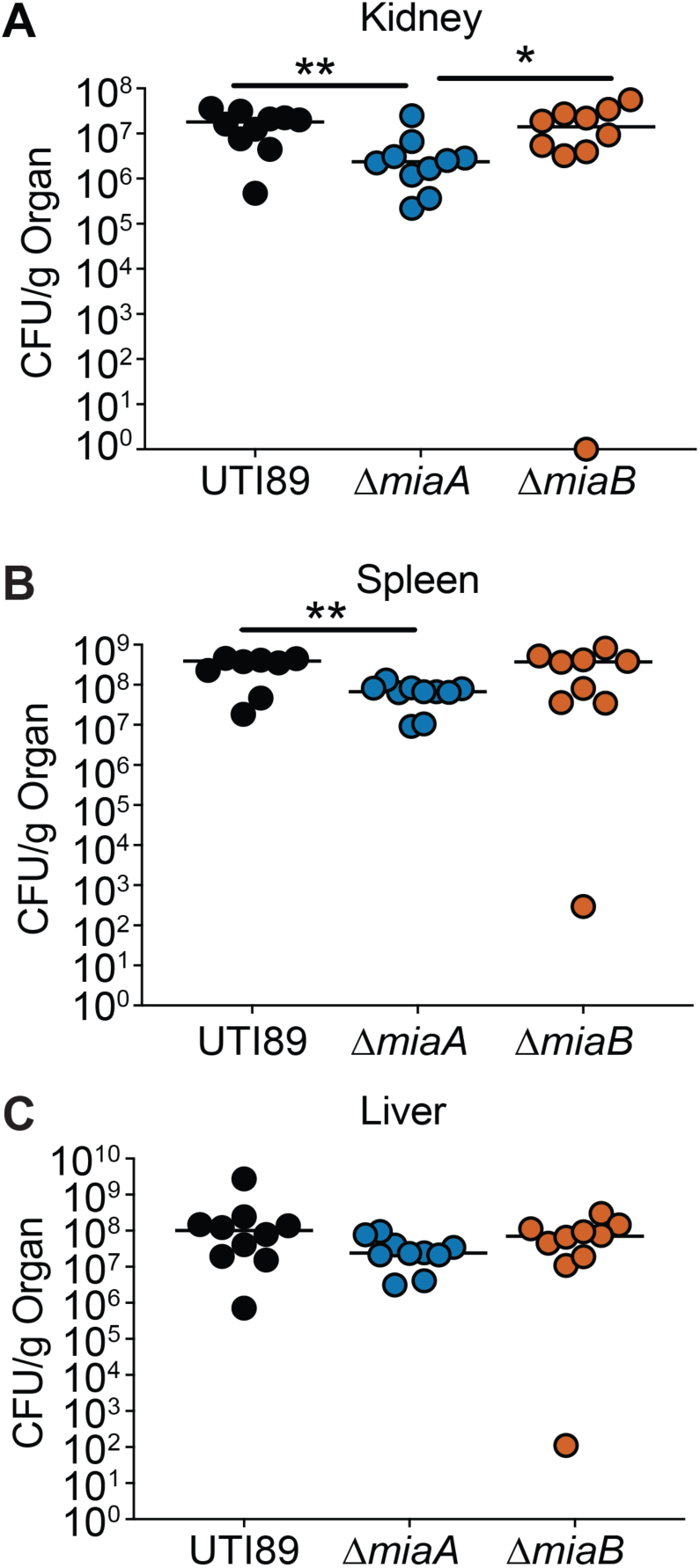
MiaA promotes ExPEC fitness in a mouse model of sepsis. Adult female C57Bl/6 mice were inoculated via i.p. injections with 10^7^-10^8^ CFU of UTI89, UTI89Δ*miaA*, or UTI89Δ*miaB* and 6 hours later bacterial titers were present in the (**A**) kidneys, (**B**) spleen, and (**C**) liver were determined by plating tissue homogenates. *, *P* < 0.05 and **, *P* > 0.01 by Mann Whitney U tests; *n* = 10 mice per group.

**Supplemental Figure S4.**
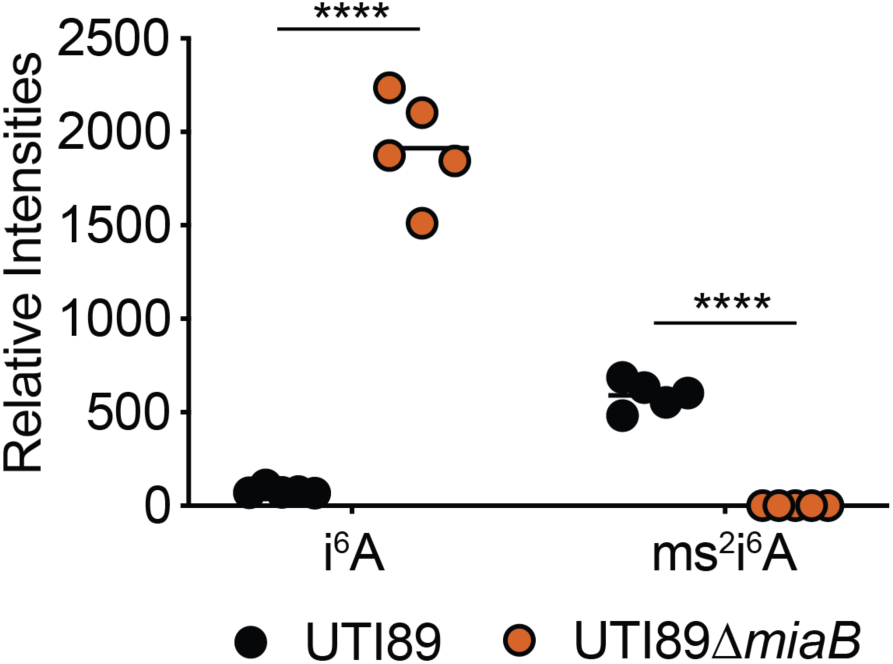
The ms^2^i^6^A modification is missing in UTI89Δ*miaB*. RNA was collected from wild-type UTI89 and UTI89Δ*miaB* after reaching an OD_600_ of 0.5 in shaking LB cultures. Relative levels of i^6^A and ms^2^i^6^A were determined by LC-MS. ****, *P* < 0.0001 as determined an unpaired *t* test; n = 5 independent replicates.

**Supplemental Figure S5.**
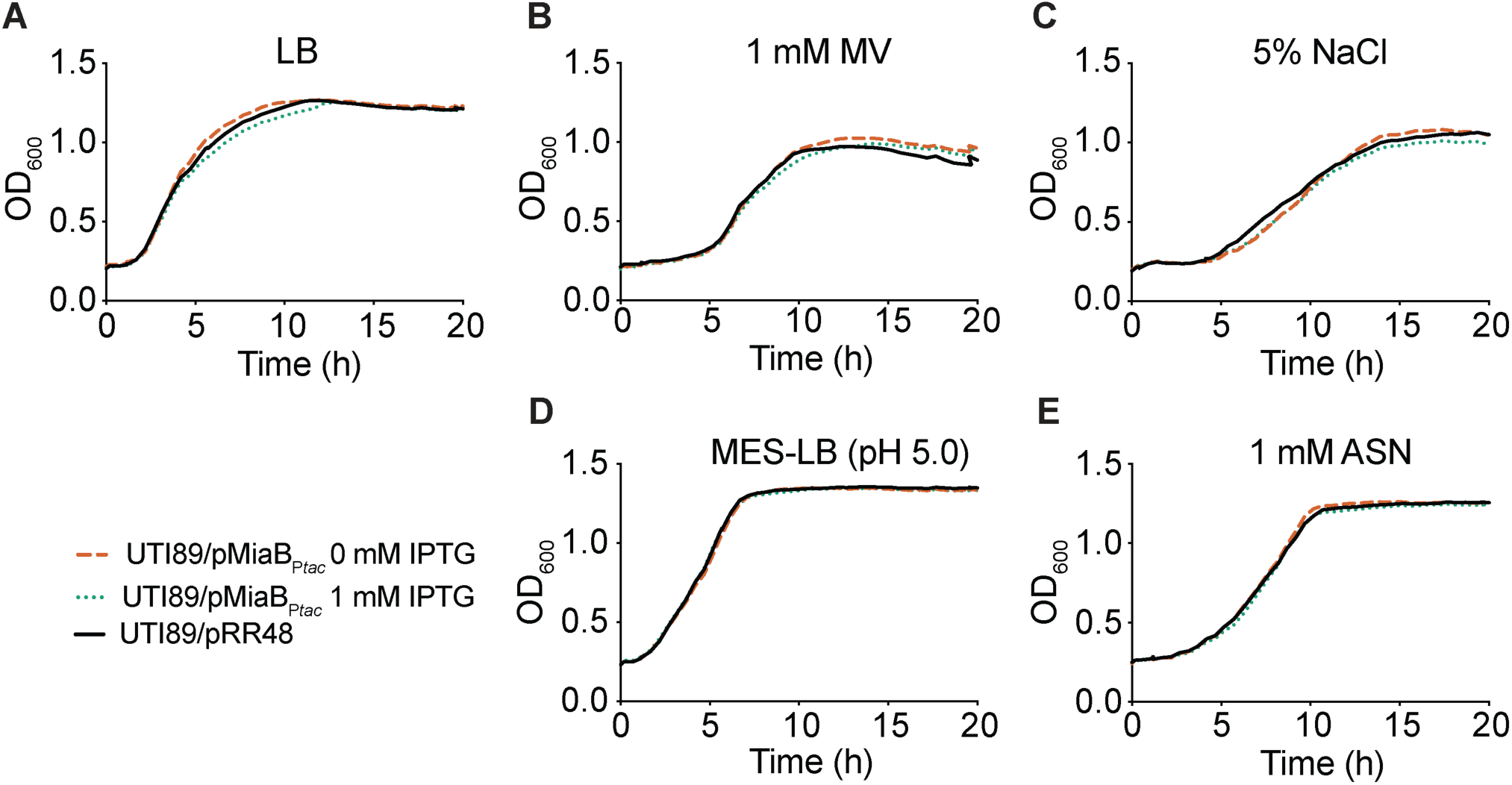
Overproduction of MiaB does not affect growth of UTI89 under stressful conditions. Graphs show growth curves of UTI89 carrying pMiaB_P*tac*_ or the control plasmid pRR48 in (**A**) LB, (**B**) 1 mM MV, (**C**) 5% NaCl, (**D**) MES-LB, and (**E**) 1 mM ASN. To overexpress MiaB, 1 mM IPTG was added to UTI89/pMiaB_P*tac*_. Data are representative of three or more independent experiments, each done in quadruplicate.

**Supplemental Figure S6.**
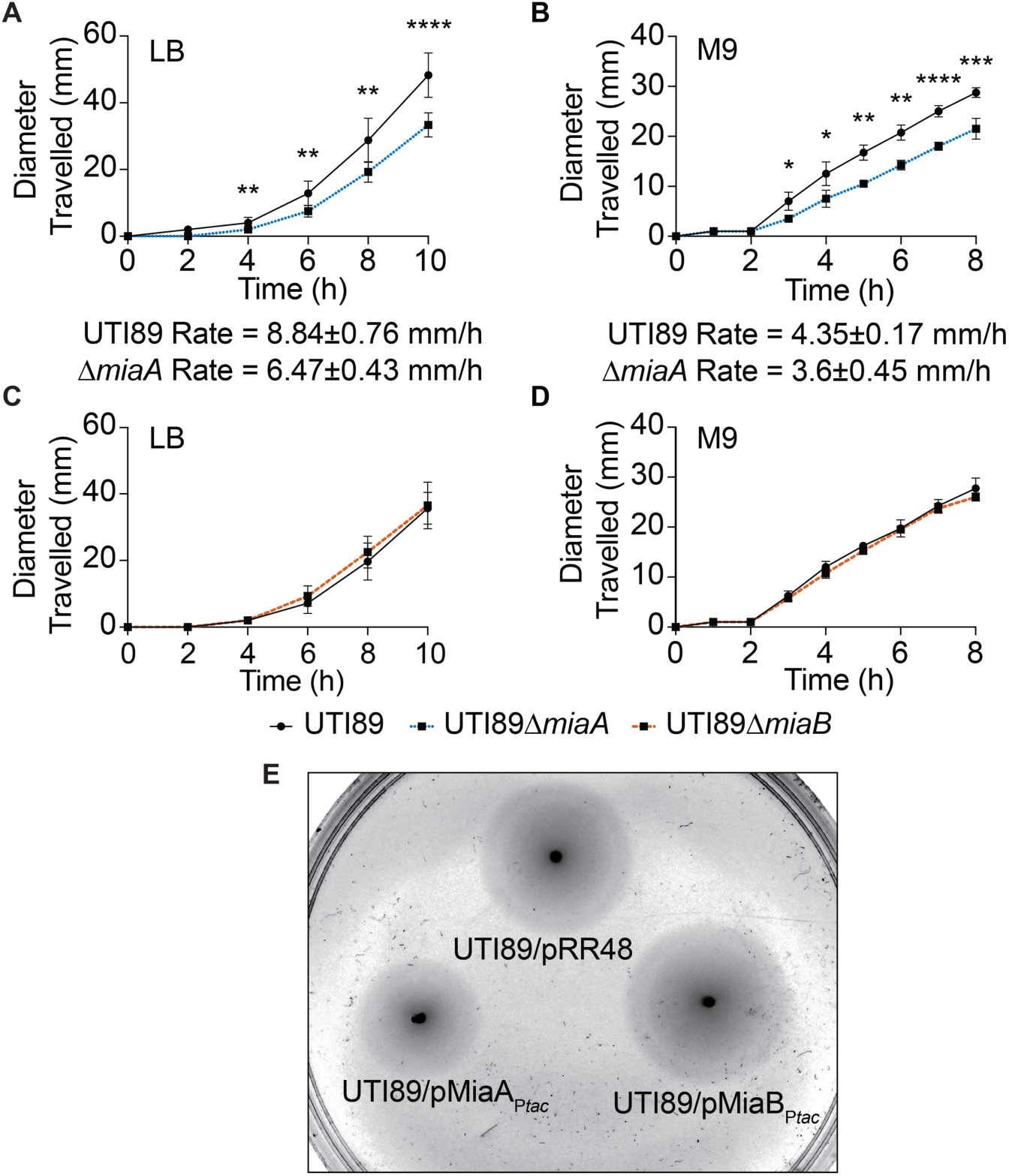
MiaA modulates ExPEC motility. (**A - D**) Graphs indicate the spread of UTI89 (black lines), UTI89Δ*miaA* (dotted blue lines), and UTI89Δ*miaB* (dashed red lines) on (A and C) LB and (B and D) M9 swim motility plates incubated at 37°C. Shown are mean values ± SD from three independent experiments done in triplicate. Swim rates (± SD) for wild-type UTI89 and UTI89Δ*miaA* on LB and M9 swim plates are indicated below the graphs in (A) and (B). *, *P* < 0.05; **; *P* < 0.01; ***, *P* < 0.001; ****, *P* < 0.0001 versus wild-type UTI89, as determined by unpaired *t* tests; n ≥ 4 independent replicates. (**E**) Representative image showing the spread of UTI89/pRR48, UTI89/pMiaB_P*tac*_, and UTI89/pMiaA_P*tac*_ 6 hours after inoculation onto an LB swim plate containing 1 mM IPTG and ampicillin.

**Supplemental Figure S7.**
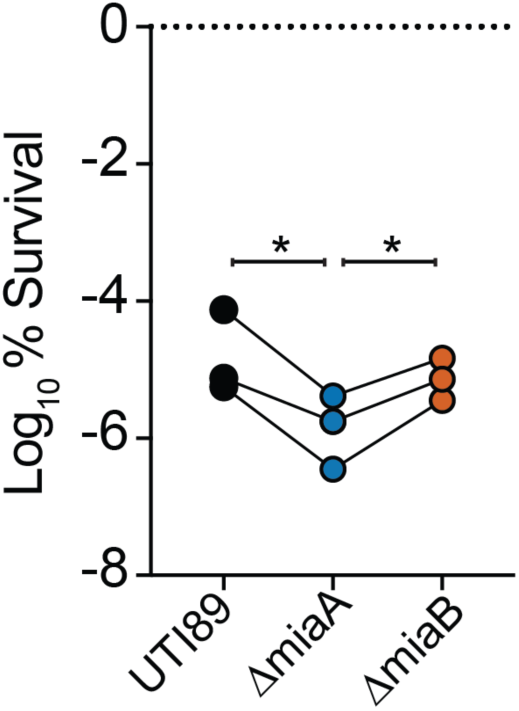
UTI89Δ*miaA* is has increased sensitivity to acid stress. After reaching mid-logarithmic growth phase in LB, wild-type UTI89, UTI89Δ*miaA*, and UTI89Δ*miaB* were exposed to acidic stress (pH 3.0) for 30 min. Following washes in PBS, surviving bacteria were enumerated by dilution plating. Titers are normalized to input. Biological replicates are connected by lines. *, *P* < 0.05 by paired *t* tests.

**Supplemental Figure S8.**
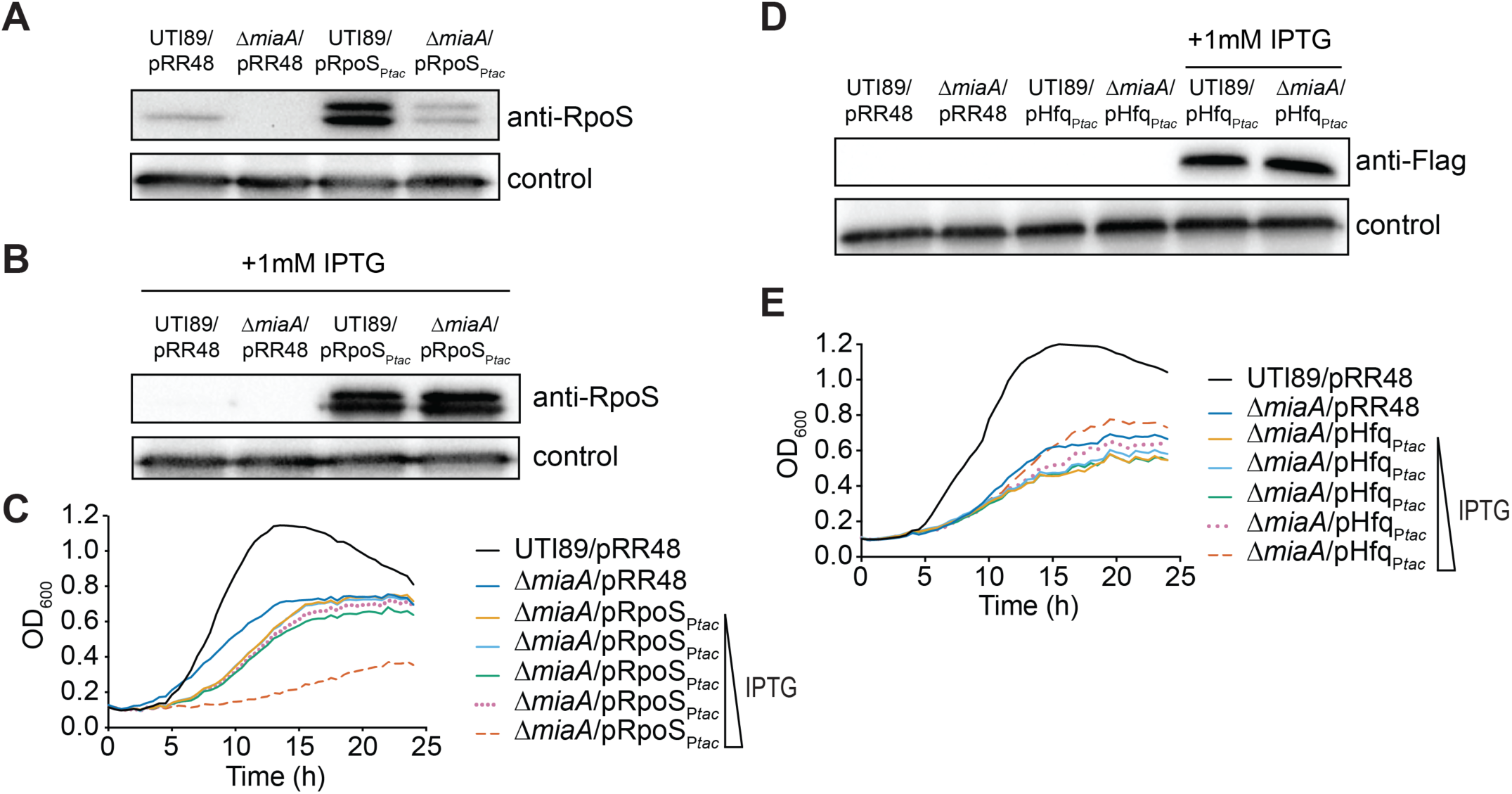
Expression of RpoS or Hfq does not rescue growth of UTI89Δ*miaA* in the presence of high salt stress. (**A - C**) Western blots of RpoS and Flag-tagged Hfq in UTI89 and UTI89Δ*miaA* carrying pRpoS_P*tac*_, pHfq_P*tac*_, or the empty vector pRR48 following growth to stationary phase in LB or LB with 1 mM IPTG, as indicated. As a loading control, blots were also probed with anti-*E. coli* antibody. A shorter exposure was used for the blot shown in (B), making the RpoS band from UTI89/pRR48 notably lighter than the one shown in (A). Blots are representative of three independent experiments. (**D and E**) Curves show growth of the UTI89 and UTI89Δ*miaA* with the empty vector pRR48 or plasmids for IPTG-inducible expression of RpoS or Flag-tagged Hfq in LB + 5% NaCl. Cultures were grown shaking at 37°C with IPTG added in ten-fold increments from 0 to 1000 μM, as indicated. Each growth curve shows the means of results from a single experiment and is representative of at least three independent experiments performed in quadruplicate.

**Supplemental Figure S9.**
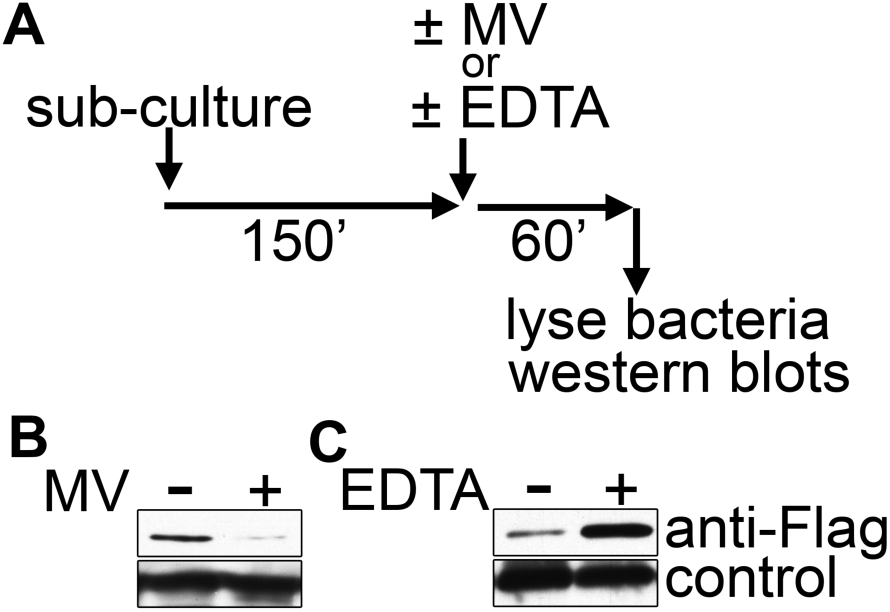
Methyl viologen and EDTA alter MiaA levels. (**A**) Top panel shows schematic of the experimental setup. UTI89/pMiaA-Flag_nat_ was diluted from overnight cultures into fresh LB and grown shaking for 2.5 hours at 37°C prior to resuspension in LB, LB + 5% NaCl, or LB + 1 mM EDTA. Incubations were continued for another hour before samples were collected and analyzed by western blots. (**B** and **C**) Blots were probed using anti-Flag (MiaA-Flag) and anti-*E. coli* (loading control, bottom) antibodies.

**Supplemental Figure S10.**
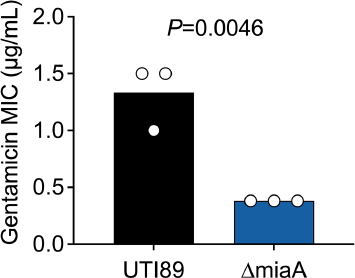
UTI89Δ*miaA* is more sensitive to gentamicin than the wildtype UTI89. Bars in graph indicate mean MIC values (± SD) determined from three independent Etests. *P*-value was determined by an unpaired Student’s *t* test.

**Supplemental Table S1.** Bacterial strains and plasmids.

| Strain or Plasmid | Description | Source or Reference |
| --- | --- | --- |
| **Strains** |  |  |
| UTI89 | UPEC reference strain, cystitis isolate | [52, 108] |
| UTI89::Kan<sup>R</sup> | UTI89 with a Kan<sup>R</sup> resistance cassette inserted at the *attTn7* site | This study |
| UTI89Δ*miaA* | UTI89 *miaA*::Cam<sup>R</sup> | This study |
| UTI89Δ*miaB* | UTI89 *miaB*::Cam<sup>R</sup> | This study |
| **Plasmids** |  |  |
| pACYC184 | Low-copy number plasmid; Tet<sup>R</sup>, Cam<sup>R</sup> | New England Biolabs |
| pBAD18 | Arabinose-inducible bacterial expression plasmid; Amp<sup>R</sup> | [109] |
| pBAD33 | Arabinose-inducible bacterial expression plasmid; Cam<sup>R</sup> | [109] |
| pKD3 | Carries FRT-flanked Cam<sup>R</sup> cassette; template for use in lambda-Red-mediated recombination | [48] |
| pKD4 | Carries FRT-flanked Kan<sup>R</sup> cassette; template for use in lambda-Red-mediated recombination | [48] |
| pKM208 | IPTG-inducible lambda Red recombinase expression plasmid; Amp<sup>R</sup> | [49] |
| pRR48 | Contains IPTG-inducible P<sub>tac</sub> promoter upstream of the MCS; Amp<sup>R</sup> | [110] |
| pHfq<sub>P<sub>tac</sub></sub> | *hfq* from UTI89 cloned with c-terminal 6xHis and FLAG tags into PstI, HindIII sites of pRR48; Amp<sup>R</sup> | This Study |
| pRpoS<sub>P<sub>tac</sub></sub> | *RpoS* cloned from UTI89 into PstI, HindIII sites of pRR48; Amp<sup>R</sup> | [62] |
| pMiaA<sub>P<sub>tac</sub></sub> | *miaA* cloned from UTI89 into PstI, KpnI sites of pRR48; Amp<sup>R</sup> | This study |
| pMiaB<sub>P<sub>tac</sub></sub> | *miaB* cloned from UTI89 into PstI, KpnI sites of pRR48; Amp<sup>R</sup> | This study |
| pMiaA<sub>nat</sub> | *miaA* plus 200 bp of flanking sequences cloned from UTI89 into the EcoR1 site of pACYC184; Tet<sup>R</sup> | This study |
| pMiaA-Flag<sub>nat</sub> | pACYC184-derived plasmid encoding MiaA with N-terminal Flag tag plus linker under control of native *miaA* promoter; Tet<sup>R</sup> | This study |
| p2Luc-HIV | Eukaryotic reporter construct with the HIV *gag-pol* frameshift region inserted between the renilla and firefly luciferase genes. The HIV linker sequence contains a 2-nucleotide insertion resulting in a stop codon located 6 codons after the start of firefly luciferase gene. The firefly gene is in a -1 frame relative to the upstream renilla gene; Amp<sup>R</sup> | Derived from [76] |
| p2Luc-HIV-IF | Control for p2Luc-HIV. The HIV *gag-pol* linker was altered to keep the renilla and firefly luciferases in-frame; Amp<sup>R</sup>. | Derived from [76] |
| p2Lucaz1 | Eukaryotic reporter construct with the Az1 frameshift region inserted between the renilla and firefly luciferase genes. The Az1 linker sequence contains a stop codon positioned in-frame so that a +1 frameshift must occur for read-through expression of firefly luciferase; Amp<sup>R</sup> | [75] |
| p2Lucaz1-IF | Control for p2Lucaz1. The Az linker was altered to keep the renilla and firefly luciferases in-frame; Amp<sup>R</sup> | [75] |
| pCWR42-CamR | Dual luciferase reporter with in-frame Az1 linker cloned from p2Lucaz1-IF into pBAD33. Has a Shine-Dalgarno sequence and is under control of the arabinose-inducible P<sub>BAD</sub> promoter. Control for pCWR43-CamR; Cam<sup>R</sup> | This study |
| pCWR42-AmpR | Dual luciferase reporter with in-frame Az1 linker cloned from p2Lucaz1-IF into pBAD18. Has a Shine-Dalgarno sequence and is under control of the arabinose-inducible P<sub>BAD</sub> promoter. Control for pCWR43-AmpR; Amp<sup>R</sup> | This study |
| pCWR43-CamR | Dual luciferase reporter with Az1 linker cloned from p2Lucaz1 into pBAD33. Has a Shine-Dalgarno sequence and is under control of the arabinose-inducible P<sub>BAD</sub> promoter; Cam<sup>R</sup> | This study |
| pCWR43-AmpR | Dual luciferase reporter with Az1 linker cloned from p2Lucaz1 into pBAD18. Has a Shine-Dalgarno sequence and is under control of the arabinose-inducible P<sub>BAD</sub> promoter; Amp<sup>R</sup> | This study |
| pCWR44-CamR | Dual luciferase reporter with in-frame HIV linker cloned from p2Luc-HIV-IF into pBAD33. Has a Shine-Dalgarno sequence and is under control of the arabinose-inducible P<sub>BAD</sub> promoter. Control for pCWR45-Cam; Cam<sup>R</sup> | This study |
| pCWR44-AmpR | Dual luciferase reporter with in-frame HIV linker cloned from p2Luc-HIV-IF into pBAD18. Has a Shine-Dalgarno sequence and is under control of the arabinose-inducible P<sub>BAD</sub> promoter. Control for pCWR45-Amp; Amp<sup>R</sup> | This study |
| pCWR45-CamR | Dual luciferase reporter with HIV linker cloned from p2Luc-HIV into pBAD33. Has a Shine-Dalgarno sequence and is under control of the arabinose-inducible P<sub>BAD</sub> promoter; Cam<sup>R</sup> | This study |
| pCWR45-AmpR | Dual luciferase reporter with HIV linker cloned from p2Luc-HIV into pBAD18. Has a Shine-Dalgarno sequence and is under control of the arabinose-inducible P<sub>BAD</sub> promoter; Cam<sup>R</sup> | This study |

**Supplemental Table S2.**
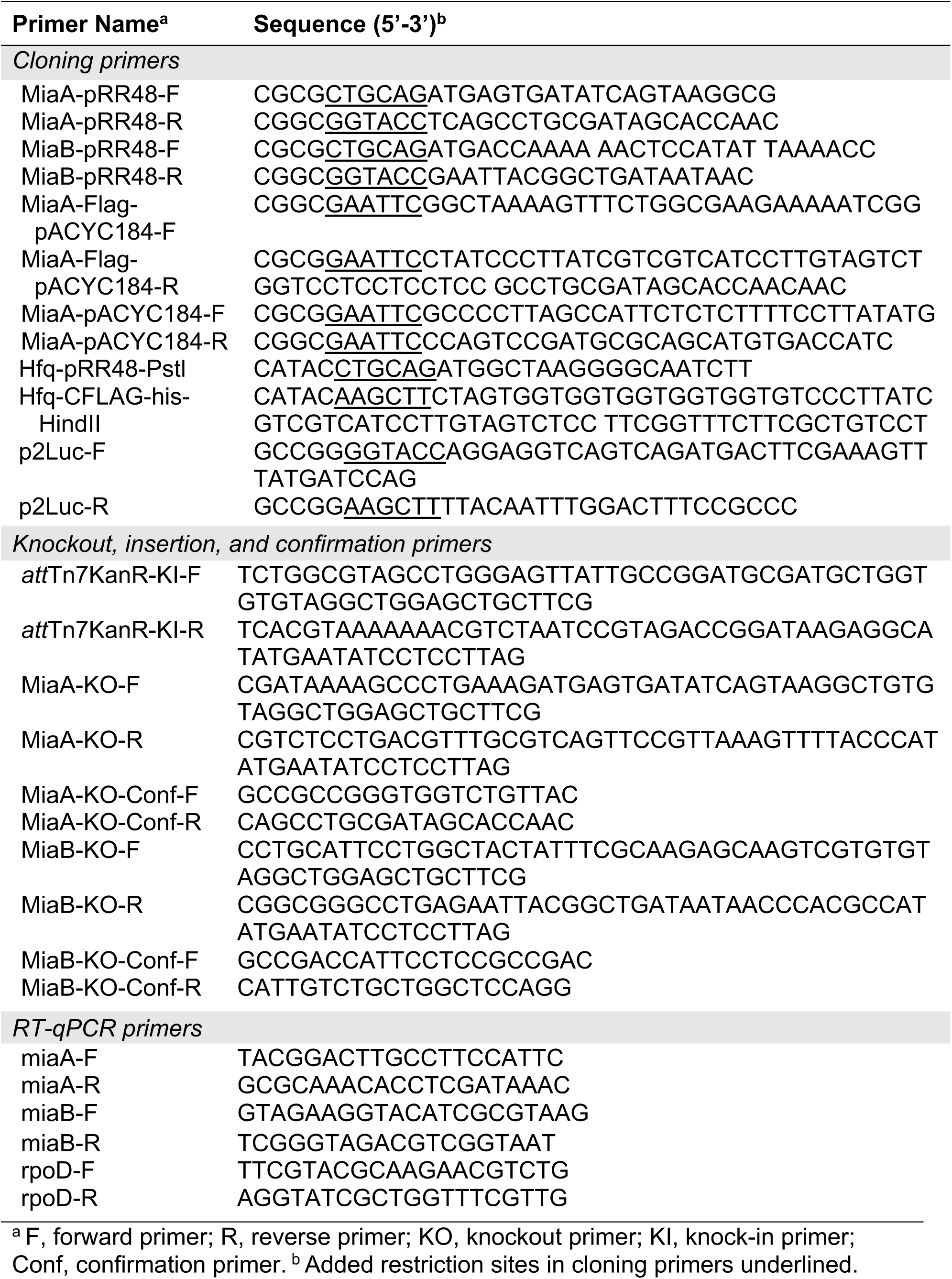
Primers used in this study.

## Notes

### Competing Interest Statement

The authors have declared no competing interest.

